# Bacterial defences interact synergistically by disrupting phage cooperation

**DOI:** 10.1101/2023.03.30.534895

**Authors:** Alice Maestri, Elizabeth Pursey, Charlotte Chong, Benoit J. Pons, Sylvain Gandon, Rafael Custodio, Matthew Chisnall, Anita Grasso, Steve Paterson, Kate Baker, Stineke van Houte, Anne Chevallereau, Edze R. Westra

**Affiliations:** Environment and Sustainability Institute, Biosciences, University of Exeter, Penryn TR10 9FE, UK; Institute of Infection, Veterinary and Ecological Sciences, University of Liverpool, Liverpool L69 3BX, UK; CEFE, CNRS, Université de Montpellier, EPHE, IRD, Montpellier 34293, France; Department of Comparative Biomedicine and Food Science, Università degli Studi di Padova, Padova 35020, Italy (current affiliation); Université Paris Cité, CNRS, INSERM, Institut Cochin, Paris F-75014, France

## Abstract

The constant arms race between bacteria and their phages has resulted in a large diversity of bacterial defence systems^1,2^, with many bacteria carrying several systems^3,4^. In response, phages often carry counter-defence genes^5–9^. If and how bacterial defence mechanisms interact to protect against phages with counter-defence genes remains unclear. Here, we report the existence of a novel defence system, coined MADS (Methylation Associated Defence System), which is located in a strongly conserved genomic defence hotspot in *Pseudomonas aeruginosa* and distributed across Gram-positive and Gram-negative bacteria. We find that the natural co-existence of MADS and a Type IE CRISPR-Cas adaptive immune system in the genome of *P. aeruginosa* SMC4386 provides synergistic levels of protection against phage DMS3, which carries an anti-CRISPR (*acr*) gene. Previous work has demonstrated that Acr-phages need to cooperate to overcome CRISPR immunity, with a first sacrificial phage causing host immunosuppression to enable successful secondary phage infections^10,11^. Modelling and experiments show that the co-existence of MADS and CRISPR-Cas provides strong and durable protection against Acr-phages by disrupting their cooperation and limiting the spread of mutants that overcome MADS. These data reveal that combining bacterial defences can robustly neutralise phage with counter-defence genes, even if each defence on its own can be readily by-passed, which is key to understanding how selection acts on defence combinations and their coevolutionary consequences.

## Introduction

Phage are important drivers of the ecology and evolution of their bacterial hosts^12^. In response to phage predation, bacteria have evolved many different defence systems, such as Restriction-Modification (RM) and CRISPR-Cas^13^. Crucially, literally dozens of previously unknown defences were discovered in recent years^2,14–19^, often aided by their clustering in defence islands^20,21^. These defence systems frequently coexist in the same genome^20–23^, which can prevent the emergence of spontaneous phage mutants that overcome host resistance. For example, the co-occurrence of a Type I BREX and Type IV RM systems prevents the emergence of epigenetic mutants that overcome BREX, since these are cleaved by the Type IV RM system^24^, and the co-existence of RM and CRISPR-Cas leads to a reduction in the frequency of spontaneous phage mutants that escape both defences as well as a higher rate of CRISPR immunity acquisition^25,26,27^. However, many phage can employ more sophisticated counter-defence mechanisms, such as anti-RM^5^, anti-CRISPR (Acr)^6^ and the more recently identified anti-CBASS^7,8^, anti-Pycsar^8^ and anti-TIR-STING^9^ proteins. We currently lack an understanding of how multi-layered bacterial defences impact the efficacy and deployment of these sophisticated phage counter-defence systems, and how this influences bacteria-phage coevolution. Previous work on the epidemiological and co-evolutionary consequences of Acr-phage infections has demonstrated that Acr are imperfect, resulting in a high proportion of failed infections of CRISPR-immune bacteria^6^. This imperfection is at least in part due to Acr production taking place after infection (i.e., Acr are not packaged into the phage capsid)^6,28^, whereas CRISPR immune complexes are already present in the cell upon infection. Because of this, Acr-phages must cooperate to overcome CRISPR immunity: the production of Acr proteins by a first sacrificial phage into a CRISPR immune host cell is necessary to enable a second Acr-phage to successfully infect the same host^10,11^. As a consequence, Acr-phages can amplify only if their density exceeds a critical threshold that supports the required frequency of secondary infections. However, our understanding of these dynamics is currently limited to the simple scenario where CRISPR-Cas immune systems are the sole resistance determinants, and it is unclear if and how this may be impacted by the presence of other defence genes in the bacterial genome.

## Results

### Bacteria drive Acr-phages extinct

We aimed to study how the Type IE CRISPR-Cas immune system of *P. aeruginosa* SMC4386 shapes the coevolutionary interactions with its phage DMS3. The CRISPR-Cas immune system of this bacterial strain carries a spacer that perfectly matches the genome of DMS3, whereas the phage carries an anti-CRISPR gene (*acrIE3*) that blocks the Type IE CRISPR-Cas system^29^. However, infection experiments of *P. aeruginosa* SMC4386 with phage DMS3*vir*, a c-repressor mutant locked in the lytic cycle, revealed that, although phages successfully adsorbed to the bacteria (Extended Data Fig. 1a), they were unable to amplify and instead rapidly went extinct, even at high initial multiplicity of infection (MOI) (Extended Data Fig. 1b). Moreover, infection experiments with the temperate phage DMS3-Gm, a mutant of phage DMS3 that carries a gentamycin resistance gene^11^ which allows the selection for lysogens, revealed that no lysogens were formed in the SMC4386 strain, while lysogens could be observed for the reference PA14 strain (Extended Data Fig. 1c). The absence of lysogen formation in SMC4386 contrasts with previous work showing efficient lysogenization on bacteria with CRISPR-Cas immunity when phages carry *acr* genes^6,11,30^.

Previous studies have shown that Acr proteins are imperfect and vary in their strength, ranging from highly effective to weak inhibitors of CRISPR-Cas^10,11^. If AcrIE3 was a weak inhibitor of CRISPR-Cas, it would explain the lack of phage amplification or lysogen formation on CRISPR immune bacteria. To test this hypothesis, we compared the efficiency of plating (EOP) of phage DMS3*vir* and a phage mutant lacking the *acrIE3* gene (referred to as DMS3*vir*Δ*acr*). While DMS3*vir* could form plaques on SMC4386, DMS3*vir*Δ*acr* could not, supporting the hypothesis that AcrIE3 efficiently blocks the Type IE CRISPR-Cas immune system of SMC4386 (Extended Data Fig. 1d). Consistent with this observation, phage DMS3*vir*Δ*acr* was able to form plaques on a CRISPR-Cas deletion mutant (SMC4386ΔCRISPR, referred to as ΔCRISPR) (Extended Data Fig. 1d). Moreover, unlike DMS3*vir*Δ*acr*, we did not detect an increase in DMS3*vir* EOP on the ΔCRISPR strain compared to the wildtype SMC4386 (SMC4386-WT) (Fig. 1a, compare yellow and black bars). Collectively, these data suggest that AcrIE3 is an effective inhibitor of the Type IE CRISPR-Cas system of *P. aeruginosa* SMC4386.

**Fig. 1.**
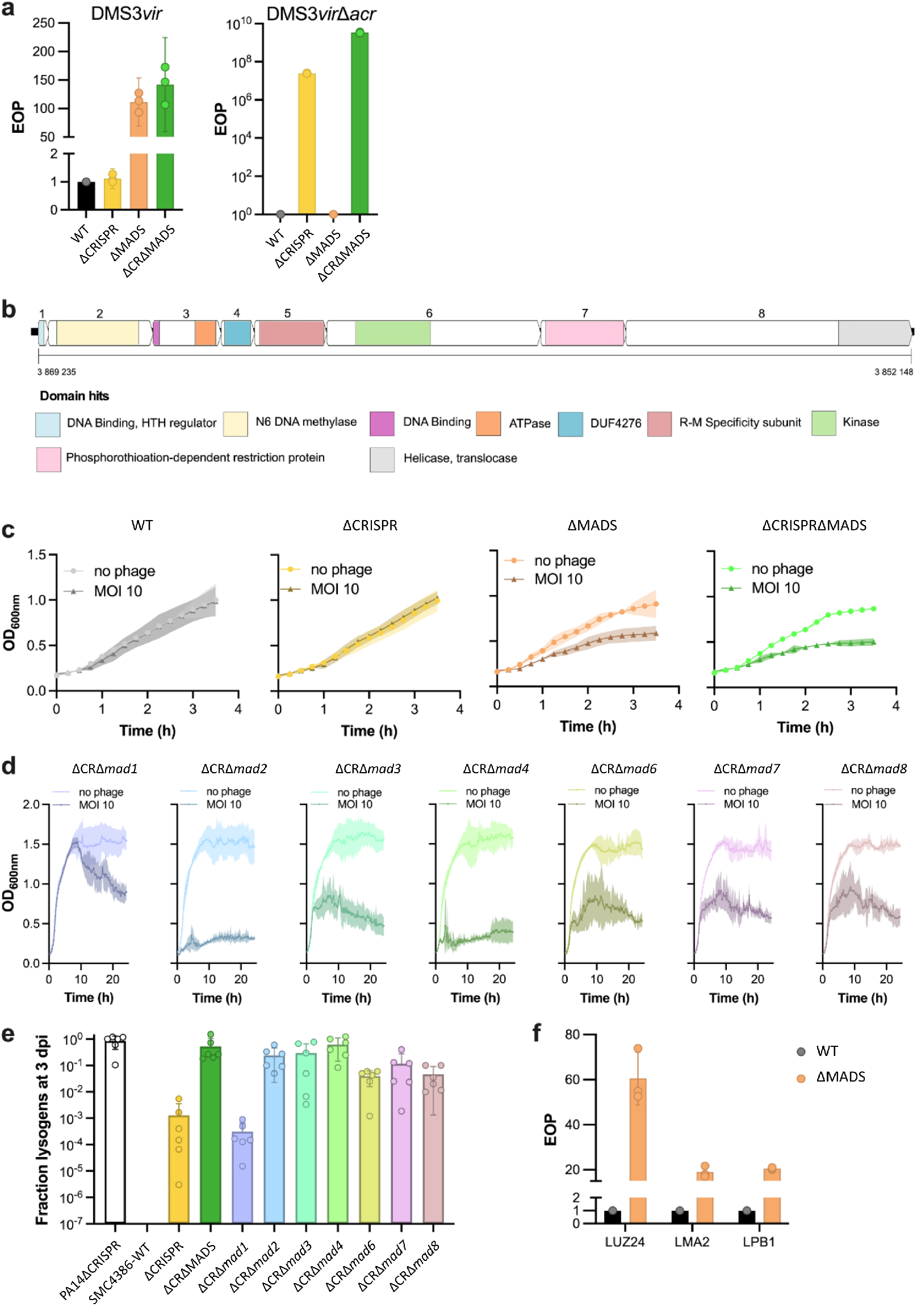
| MADS protects bacteria against phages. **a,** Efficiency of plating (EOP) of phage DMS3*vir* or DMS3*vir*Δ*acr* on strains SMC4386-WT, ΔCRISPR, ΔMADS and ΔCRISPRΔMADS. **b**, MADS is an 8-gene operon. Numbers in the top arrows correspond to gene numbering (*mad1*-*mad8*). Colours in bottom arrows indicate domain hits predicted with HHpred analyses and the scale indicates the position of the *mad* operon on the SMC4386 genome assembled as single chromosome (in bp). **c**, **d**, Bacterial growth curves of WT and mutant strains in absence or presence of phage DMS3*vir* (with multiplicity of infection, MOI, of 10). Single *mad* gene deletions are all in a ΔCRISPR background (indicated as ΔCRΔ*madX,* with X the gene number). **e**, Fraction of the bacterial population carrying the DMS3 prophage (lysogens) in WT and mutant SMC4386 backgrounds, as indicated. ΔCR is an abbreviation for ΔCRISPR. **f**, EOP of phages LUZ24, LMA2 and LBP1 on SMC4386-WT and the isogenic ΔMADS strain. For each panel, values for individual replicates are shown (3 replicates in panels a and f, 4 replicates in c and d and 6 replicates in panel e) as well as mean values. Errors bars (panels a, e, f) and shaded areas (panels c, d) show 95% confidence intervals (c.i.).

### A novel defence system coined MADS

Based on these observations, we hypothesised that *P. aeruginosa* SMC4386 may carry additional defence systems that limit DMS3 infectivity. To test this prediction, we generated a transposon (Tn) mutant library of SMC4386-WT, which we then infected with DMS3-Gm. Interestingly, lysogen formation was observed in the SMC4386-Tn mutant library (Extended Data Fig. 2c, yellow line), but not in the SMC4386-WT strain (Extended Data Fig. 1c). PCR analysis of the DMS3 c-repressor gene confirmed that 83.6% of the Gm-resistant clones within the SMC4386-Tn mutant population were lysogens, and not spontaneous Gm-resistant mutants. Sanger sequencing of these lysogens revealed that most of the Tn insertions were located in *cas* genes (Extended Data Fig. 2b), supporting the idea that Acr proteins are imperfect and that CRISPR-Cas plays a role in blocking DMS3 lysogen formation, despite the phage carrying an *acrIE3* gene. To test the hypothesis that additional genes, beyond CRISPR-Cas, are involved in defence against phage DMS3, we next generated a transposon mutant library of the ΔCRISPR strain. Using the same method, we identified multiple Tn insertions in a restricted genomic region containing a predicted operon of 8 genes^31,32^ (Extended Data Fig. 2c). This putative operon did not contain a known defence system based on PADLOC and DefenseFinder analyses^3,33^ (Supplementary Table 1). However, PADLOC identified 3 genes in this operon as putative components of DNA-modification systems: a kinase, a specificity subunit and a methylase (Supplementary Table 1). Therefore, we hypothesised that this operon might encode a defence system that has not been previously described, which we coined MADS (Methylation Associated Defence System) (Fig. 1b). Interestingly, this putative new 8-gene system contains predicted domains that are also found in other bacterial defence systems (see Supplementary Notes), supporting a role in defence against mobile genetic elements (MGE) (Fig. 1b). To experimentally test this hypothesis, we knocked out ∼22kb of the genome encompassing this operon (referred to as ΔMADS). Plaque assays with phage DMS3*vir* showed a 100-fold increase in the EOP on ΔMADS compared to the WT strain (Fig. 1a, orange bar). Likewise, the EOP of DMS3*virΔacr* increases by approximately 100-fold on a double mutant ΔMADSΔCRISPR compared to ΔCRISPR (Fig. 1a, compare yellow and green bars). The observed differences in phage infectivity were reflected in the suppression of bacterial growth in the presence of phages. When MADS is not present (ΔCRISPRΔMADS and ΔMADS strains), bacteria displayed normal growth in the absence of phage DMS3*vir* but reduced growth in its presence (Fig. 1c, orange and green curves), whereas in the presence of MADS (WT and ΔCRISPR strains) bacteria grew equally well in the presence and absence of phage at a MOI of 10 (Fig. 1c, black and yellow curves).

Having established that this operon encodes a novel defence system, we next wanted to test the implication of each gene in this phenotype (*mad1-8*, Fig. 1b). We therefore generated knock-out mutants of each individual gene and measured how this impacted the phage resistance. Interestingly, despite repeated attempts, we were unable to delete *mad5*, unless also *mad2* or *mad3* were deleted, suggesting that deletion of *mad5* alone is lethal. Infection experiments showed that individual gene deletions of *mad2-4* and *mad6-8* all resulted in reduced bacterial growth in the presence of phage DMS3*vir*, whereas deletion of *mad1* had limited effect (Fig. 1d). Moreover, deletion of either the full system or each of the individual *mad* genes resulted in a 30- to 400-fold increase in phage DMS3 lysogen formation in mutant backgrounds relative to the ΔCRISPR strain, except for *mad1* (Fig. 1e). To understand the range of protection offered by MADS, we carried out infection assays with diverse phages, including temperate phage LPB1(*Casadabanvirus*) and lytic phages LMA2 (*Pbunavirus*) and LUZ24 (*Bruynoghevirus*). We observed a similar increase (>10-fold) in the EOP of all phages in ΔMADS strain relative to the SMC4386-WT (Fig. 1f). Based on these data, we conclude that the MADS system encodes a novel defence system that is active against at least 4 different phages, with genes *mad2-4* and *mad6-8* being essential for phage resistance.

### Distant bacterial classes carry MADS

To determine how widespread and conserved MADS is, we built a MacSyFinder model^34^ and searched through all bacterial and archaeal genomes in the RefSeq database. We identified 422 MADS in 100 different species belonging to *Alpha, Beta, Gamma* and *Deltaproteobacteria*, *Actinobacteria* and *Nostocales* (Extended Data Fig. 3a). The *mad1-8* operon was most commonly found in the genera *Escherichia* (n=113, 27% of systems detected), *Pseudomonas* (n=71, 17%), *Ralstonia* (n=51, 12%), *Streptomyces* (n=33, 8%), *Klebsiella* (n=31, 7%) and *Vibrio* (n=15, 4%) (Extended Data Fig. 3b). This analysis also revealed varying levels of completeness of MADS operon, with some genes missing or being detected less frequently in some bacterial genera, suggesting the existence of multiple subtypes of MADS (Extended Data Fig. 3c,d). The majority of operons identified in the search contained 8 or more genes, including multiple hits to the same gene type, possibly reflecting gene duplication events (Extended Data Fig. 3e).

### MADS is located in a defence hotspot

A deeper examination of the *P. aeruginosa* genomes containing MADS revealed that the complete operon was detected in approximately 1% (66/6103) of the genomes analysed. Interestingly, manual inspection of these 66 genomes led to the observation that in at least 51/66 (77%), the MADS operon was located within a genomic region with conserved gene boundaries: *pheT*, a phenylalanine tRNA ligase subunit beta (MPAO1_RS11510) and a histidine kinase (MPAO1_RS11445). An extended search encompassing all the currently available complete *P. aeruginosa* genomes (n= 454) revealed a highly conserved hotspot for *P. aeruginosa* defence systems at this location in the genome (Fig. 2a). The gene boundaries were found in 97% (444/454) of the genomes analysed (Supplementary Table 2), delimiting genomic islands with sizes ranging from 9.5 kb to 2402.9 kb, with an average length of 64.3 kb. For the 444 genomes containing the island, PADLOC^33^ was used to annotate defence systems both on the extracted islands (Supplementary Table 3) and on the remainder of the genome (Supplementary Table 3). Analysing the number of defence systems detected by PADLOC per unit of genomic length (in kb) confirmed a statistically significant enrichment for defence systems on this island, with a median of 0.1 systems/kb in the defence island and 0.003 systems/kb in the remainder of the genome (Fig. 2b, Wilcoxon’s signed rank test, p<0.05). Further analysis revealed that this island contains at least one defence system in 81% (358/444) and two or more defence systems in 78% (346/444) of all genomes analysed, with a total of 46 different known defence systems, as well as potential novel or incomplete defence systems (Extended Data Fig.4a and Supplementary Notes). Manual inspection of such systems revealed a putative novel defence system related to the MADS system that we coined ‘MADS-like’: this is a 11-gene system (numbered *madl1* to *madl11*), which was found in 10/444 islands (Fig. 2c, Extended Data Fig.4b and Supplementary Notes). Interestingly, both MADS and the MADS-like are predicted to encode a N6-Methyltransferase (Fig. 2c and Supplementary Notes), which suggests that self/non-self discrimination relies on a methylation-based mechanism, a common strategy employed by many bacterial innate immune systems, such as Restriction-Modification, BREX and DISARM^35^.

**Fig. 2.**
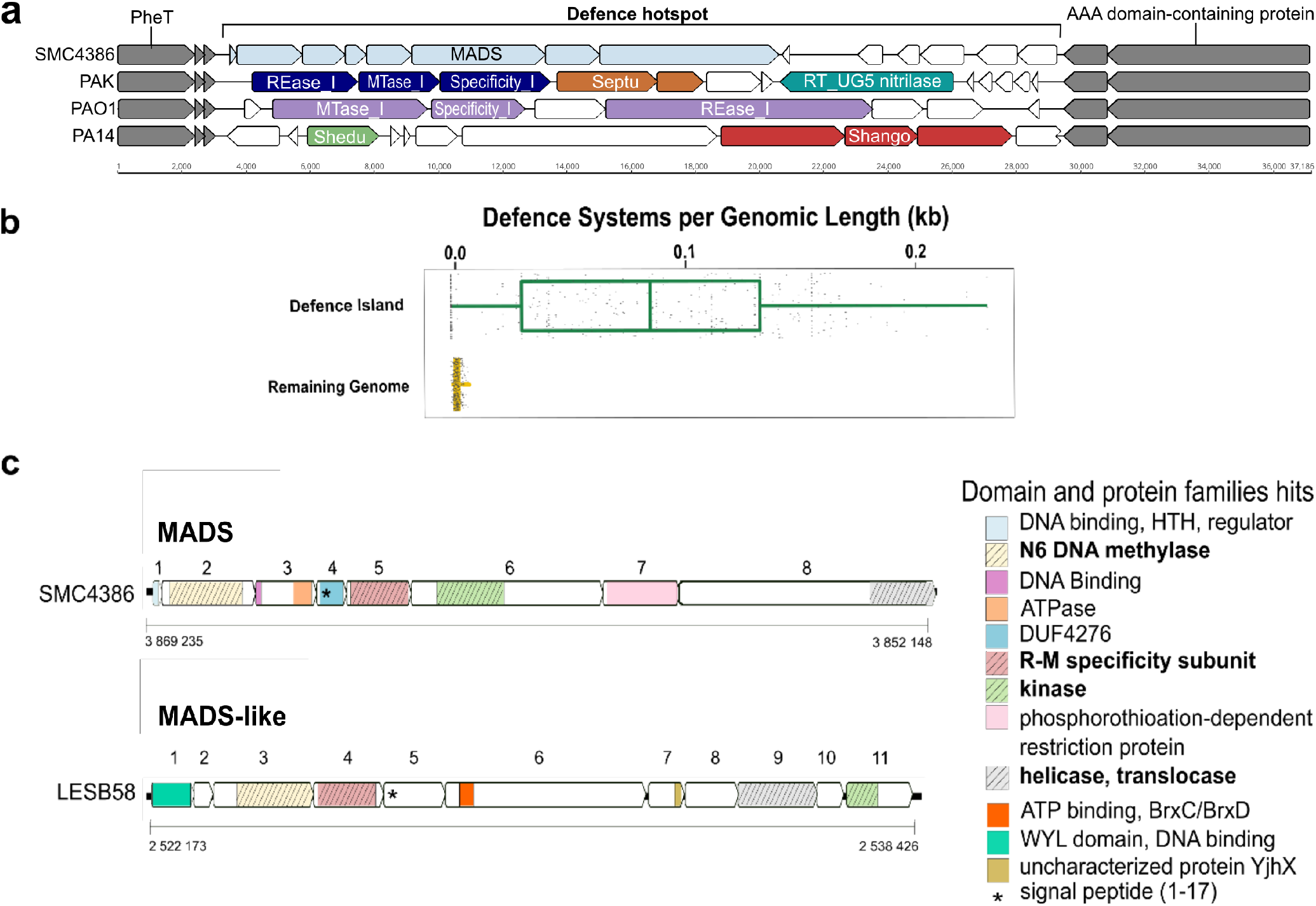
| MADS is located in an integration hotspot for bacterial defence systems. **a,** Composition of the genomic island in strains SMC4386, PAK, PAO1 and PA14. Conserved genes forming the boundaries of the island are indicated in grey. Coloured arrows indicate genes forming indicated defence systems. **b**, Number of defence systems per genomic length (kb) in the defence island compared to the remaining genome. Boxplots display the interquartile range, median and maximum defence systems per genomic length (kb) in the defence island and the remaining genome. Each grey dot represents density values for each isolate. **c**, Comparison of the MADS and MADS-like putative defence systems. Predicted protein domains (HHpred) are indicated with coloured boxes. Protein domains that are common between MADS and MADS-like are indicated in bold characters.

### Epigenetic phage mutants escape MADS

To test whether MADS uses epigenetic modification for self/non-self discrimination, we co-cultured phage DMS3*vir* and SMC4386ΔCRISPR, and monitored whether phages could evolve to overcome MADS. When the ΔCRISPR strain was infected with phage DMS3*vir*, we noticed that in some of the replicates, the phage population amplified after 2 days of infection (Fig. 3a, yellow circles with black outline). This indicates that in the absence of CRISPR-Cas, some phages evolved to overcome MADS. Interestingly, such ‘escape’ phages did not emerge during infection of the WT strain, suggesting a synergistic effect of CRISPR-Cas and MADS that prevents the evolutionary emergence of phage escape variants (Fig. 3a, black lines). The phages that evolved to overcome MADS were isolated and their infectivity was measured on different SMC4386 mutant backgrounds. This test revealed that they formed plaques with equal efficiency on all genomic backgrounds (WT, ΔCRISPR, ΔMADS and ΔCRISPRΔMADS strains, Fig 3b). To test whether the modification which allowed these escape phages to overcome MADS was epigenetic, we next amplified these phages successively on PA14ΔCRISPR (which lacks MADS) and on SMC8346-WT strains. We predicted that during amplification in the PA14 background, escape phages would lose the epigenetic modification that underpins their ability to bypass MADS, and hence lose their ability to infect SMC4386-WT in a successive passage. Consistent with this hypothesis, phages that replicated on PA14ΔCRISPR had a decreased infectivity when they were next passaged on the SMC4386-WT strain (Fig. 3c, PA14→SMC), whereas this was not the case if they replicated on the SMC4386-WT strain in the previous round of infection (Fig 3c, SMC→PA14) or if they replicated on the same strain in the previous round of infection (Fig. 3c, SMC→SMC, PA14→PA14).

**Fig. 3.**
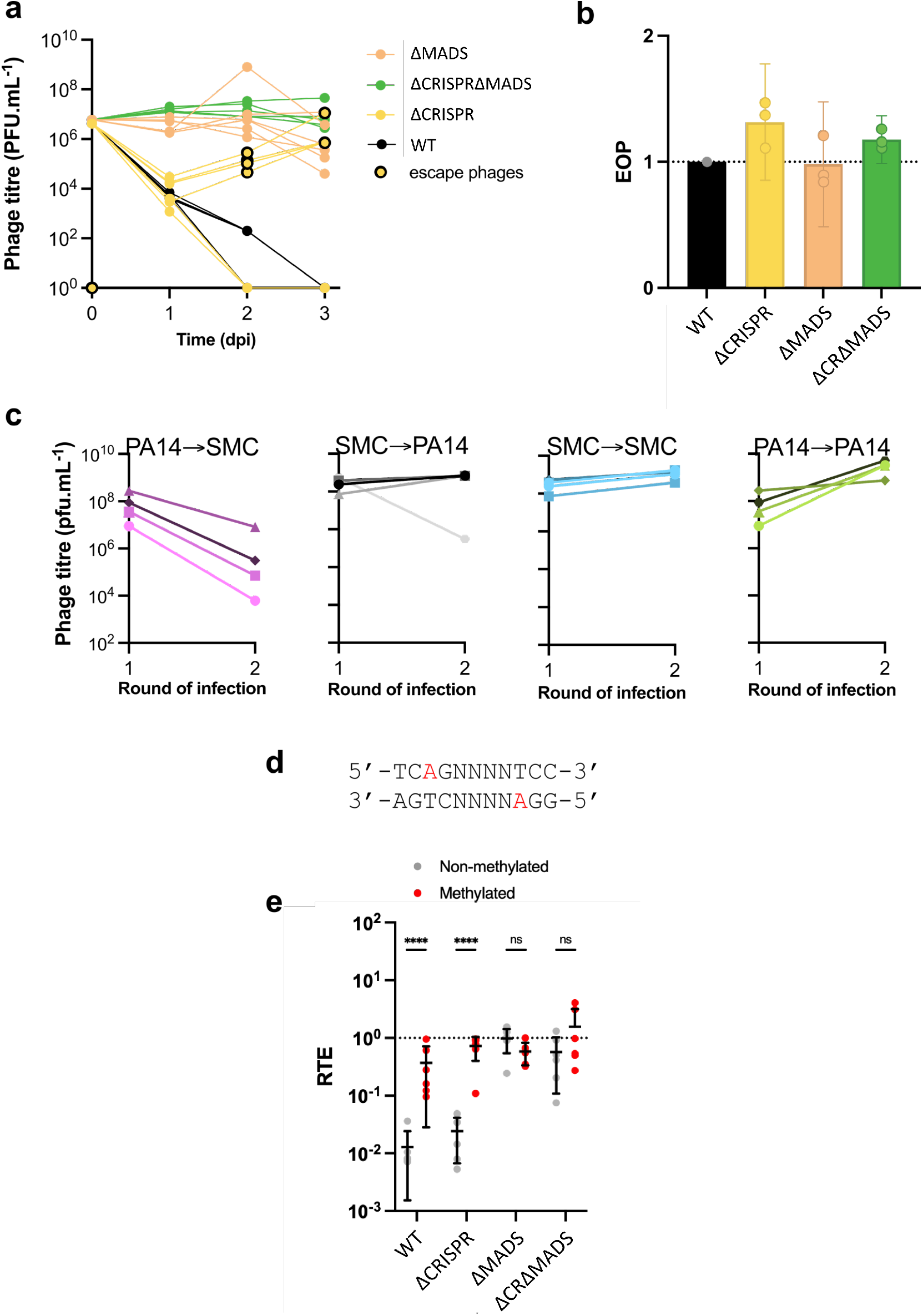
| Phages escape MADS through epigenetic modification. **a,** Titre of phage DMS3*vir* over 3 days post infection (MOI 0,1) of strains SMC8386-WT, ΔCRISPR, ΔMADS or ΔCRISPRΔMADS. Yellow circles with black outline indicate replicates from which DMS3*vir* escape phage mutants were isolated. Data are shown for 6 individual replicates. **b,** EOP of DMS3*vir* escape mutants (isolated in the experiment shown in panel a) measured on indicated strains. Individual and mean data are shown for 3 individual replicates. Error bars show 95% confidence intervals (c.i.). **c**, Amplification patterns of DMS3*vir* escape mutants when successively passaged on PA14ΔCRISPR and SMC4386-WT, with the order of passaging indicated on top of the graphs. Phage titre was assessed after each round of amplification using spot assay on PA14ΔCRISPR. Each panel shows data obtained with four independent escape phages (isolated in the experiment shown in panel a). **d**, Sequence of the genomic site that is modified by the MADS system. Adenosines that are methylated at positions 3 and 9 on the positive and negative strands (respectively) are highlighted in red. **e**, Transformation efficiency for methylated (red) or unmethylated (grey) plasmid carrying a 5’-TCAGNNNNTCC-3’ sequence, relative to a plasmid lacking this sequence.

To corroborate these phenotypic analyses and to identify the epigenetic modification site, we carried out PacBio sequencing of the escape phages and the ancestral DMS3*vir* phage. Sequencing results revealed that the motif 5’-TCAGNNNNTCC-3’ has m6A modifications at positions 3 and 9 on the positive and negative strands, respectively, in the escape phage, but not in the ancestral phage genome (Fig. 3d). The phage has 7 such motifs, all of which were methylated in the escape phages. Moreover, PacBio sequencing of the bacterial DNA revealed that the same motifs were methylated on the bacterial chromosome in the SMC4386-WT and ΔCRISPR strains, whereas these sites were not methylated in the ΔCRISPRΔMADS, nor in the ΔCRISPRΔ*mad2* strain, which encodes the predicted methyltransferase. The combination of phenotypic and sequencing data therefore supports the hypothesis that MADS uses a methylation-based self/non-self discrimination mechanism. To further corroborate the hypothesis that MADS provides resistance against MGE with unmethylated 5’-TCAGNNNNTCC-3’ sequence, but not against MGE with a methylated sequence, we measured the transformation efficiency of methylated and unmethylated plasmids with the target sequence relative to a plasmid without this sequence. Consistent with our hypothesis, a methylated plasmid carrying a 5’-TCAGCGCGTCC -3’ sequence had a 100-fold increased relative transformation efficiency (RTE) compared to the non-methylated plasmid when recipient cells carried MADS (SMC4386-WT or SMC4386ΔCRISPR, Fig. 3e), while no differences could be observed when strains lacked MADS (ΔMADS or ΔCRISPRΔMADS). Overall, our data show that recognition of the unmethylated 5’-TCAGCGCGTCC -3’ sequence by MADS is required for mediating resistance against infections by MGE.

### CRISPR-Cas and MADS act in synergy

Given that the DMS3*vir* phage has already evolved to overcome the Type IE CRISPR-Cas system of the host through the *acrIE3* gene, and that it can readily evolve to overcome MADS through the acquisition of epigenetic modifications, it is unclear why the phage is unable to persist when bacteria carry both defences (Fig. 3a, black lines). To explore the reasons why this may be the case, we developed a mathematical model (see Extended Data Fig.5a and Methods for a detailed description of the model) which builds on a previous model that considered the conditions for phage amplification in the absence of phage evolution, when bacteria have CRISPR-Cas immunity and phages carry *acr* genes^10^. This previous model assumes that protection against CRISPR immunity provided by Acr proteins is imperfect, with the probability of a successful infection depending both on the level of host resistance (ρ) and the efficacy of the Acr (ϕ). The model further assumes that during failed infections, some Acr proteins are produced and induce an immunosuppressed state in the surviving host, which reverts back to an immunocompetent state with rate γ^10^. In order to understand how the combination of adaptive and innate defences impacts phage persistence and evolution, we modified this model to include an additional defence system that prevents phage replication with efficacy *m*, which phage mutants can overcome through the acquisition of epigenetic modification. As we showed previously^10^, if *m*=0 (i.e. CRISPR-Cas is the only system that provides defence), the ability of phages with *acr* genes to amplify on CRISPR-Cas immune bacteria is density-dependent, with a critical threshold in phage density required for phage amplification (Fig. 4a, dark green line and ref^10^). However, when bacteria have both CRISPR and an additional innate immune system (i.e. *m*>0), the model predicts that the critical threshold density required for phage amplification increases towards higher phage densities (Fig 4a). This is because CRISPR-immunosuppressed cells cannot be exploited as efficiently during a secondary phage infection due to the activity of the innate immune system (such as MADS). When we examine the evolution of escape phages from the innate immune system using the same model, we find that the ability of escape mutants to amplify depends on the levels CRISPR-Cas immunity, with the threshold phage density required for phage amplification shifting to higher densities as the levels of CRISPR immunity increase (increasing values of ρ in the model, Fig. 4b). This is because infections of bacteria that are more resistant to immunosuppression limit the spread of epigenetic phage mutants in the population.

**Fig. 4.**
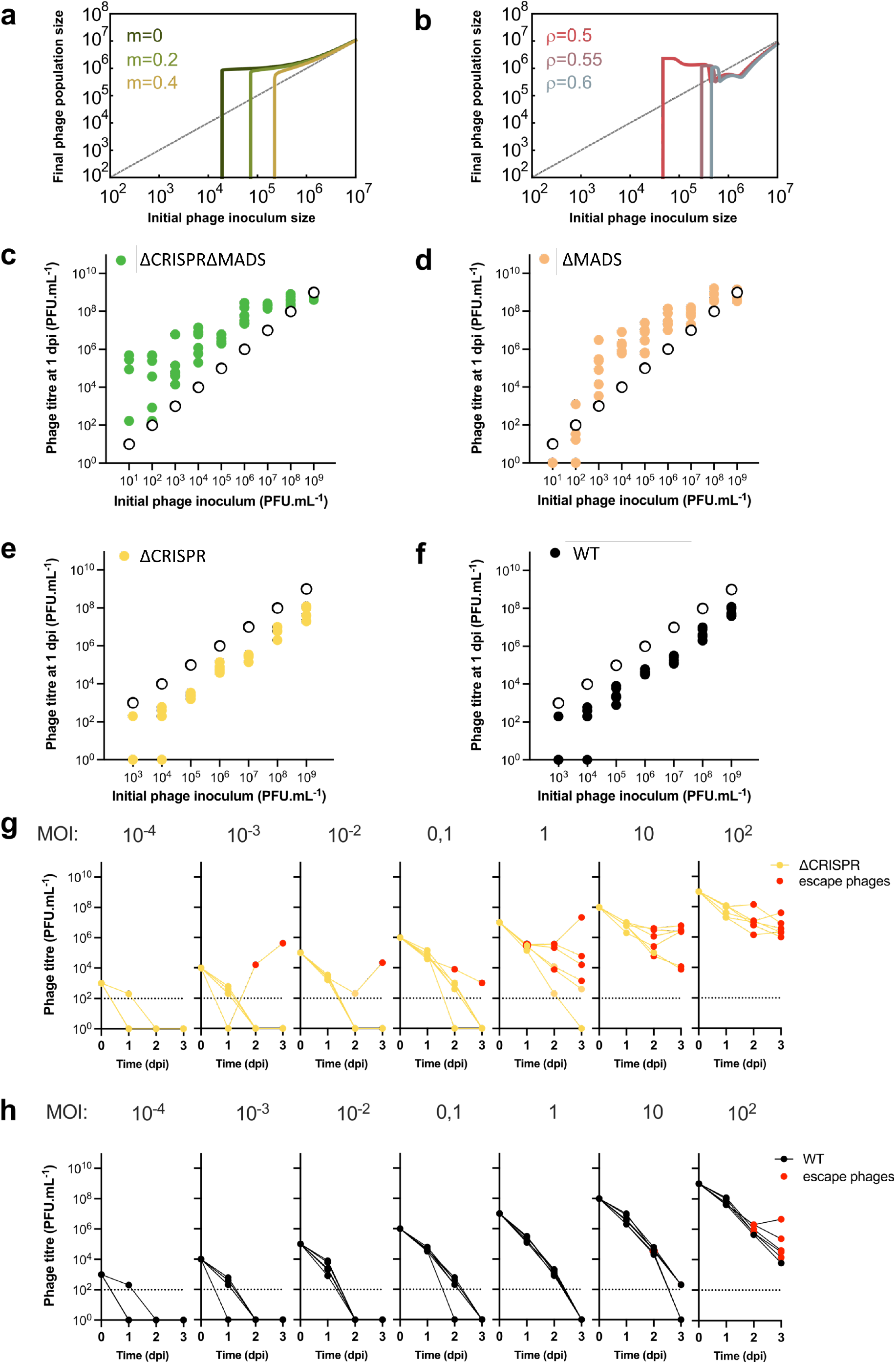
| Synergy between CRISPR-Cas and MADS. **a**, Final phage density for different values of MADS efficacy of resistance (*m*) when *ρ* = 0.6. **b**, Final phage density for different values of CRISPR efficacy of resistance (*ρ*). In both plots the initial density of resistant cells is *K*/2. Other parameter values: *m* = 0.8, *r* = 1, *γ* = 20, *ϕ* = 0.4, *B* = 5, *a* = 0.001, *d* = 0.001, *λ* = 1, *μ* = 0.0001, *K* = 10^7^. **c-f**, DMS3*vir* titre at 1 dpi upon infection of strains that (**c, d**) lack or (**e, f**) carry MADS. Empty circles show initial phage titres and coloured dots show titre at 1 dpi. **g-h,** Titre of phage DMS3*vir* over 3 days upon infection of strains (**g**) SMC4386ΔCRISPR and (**h**) SMC4386-WT. Each graph shows infection experiments starting with initial phage concentrations varying from 10^3^ to 10^9^ PFU/mL (with 10-fold increments) corresponding to initial MOIs ranging from 10^-4^ to 10^2^. Red circles highlight replicates where escape phages could be detected. All panels show individual data for 6 independent replicates.

To test the model predictions, we first validated that DMS3*vir* infection causes immunosuppression of the SMC4386 Type IE CRISPR-Cas system, as this assumption was based on studies using the *P. aeruginosa* PA14 model that carries a Type IF CRISPR-Cas system^10,11^. To test this, we briefly pre-exposed either the SMC4386-WT or the ΔMADS strain to phage DMS3*vir*, washed away all phages, followed by transformation of a CRISPR-targeted or a non-targeted plasmid^36^. We found that the transformation efficiency of the targeted plasmid, relative to the non-targeted plasmid, increased following pre-exposure to phage DMS3*vir*, but not DMS3*vir*Δ*acrIE3.* This demonstrates that failed infections cause bacteria to become immunosuppressed when the phage carries the *acrIE3* gene (Extended Data Fig.5b). Moreover, a similar increase in the relative transformation efficiency of the targeted plasmid was observed when bacteria encode only one (the ΔMADS strain) or both defence systems (SMC4386-WT) (Extended Data Fig.5b). Hence, these data validate the model assumption that phages carrying *acrIE3* are imperfect in their ability to by-pass CRISPR-Cas, and that failed infections cause cells to enter into an immunosuppressed state.

Next, we performed infection assays over a broad range of initial phage inoculi to test that (i) phage amplification requires higher initial phage densities when both defences are present and (ii) that, in the presence of CRISPR-Cas, escape phage emergence is suppressed compared to the situation where bacteria only carry MADS. This assay revealed consistent phage amplification one day post infection (dpi), regardless of the initial phage inoculum, when bacteria lack both defence systems (i.e., ΔCRISPRΔMADS) (Fig. 4c). However, when bacteria only carry the CRISPR-Cas immune system (i.e., ΔMADS), phage extinctions were observed at 1 dpi in all replicates at an initial phage inoculum of 10 PFU/mL and in 5 out of 6 replicates at an initial phage inoculum of 100 PFU/mL (Fig. 4d), whereas phage amplification was observed at higher initial phage densities, consistent with previous observations that Acr-phage amplification on CRISPR-immune bacteria depends on the initial phage density^10,11^. In contrast, when bacteria carry only MADS (i.e., ΔCRISPR) we never observed phage amplification at 1 dpi, even at the highest phage density (10^9^ PFU/mL) (Fig. 4e). Crucially, when bacteria carry both CRISPR-Cas and MADS (SMC4386-WT), we again could not observe amplification at 1 dpi across all of the initial phage inoculi (Fig. 4f). This demonstrates that at 1 dpi, phages have not yet evolved to overcome MADS, explaining why Acr-phages cannot amplify on WT bacteria.

However, we frequently observed later in the experiment that the phage population started to recover, but only when bacteria carried a stand-alone MADS (ΔCRISPR) (Fig. 4g). Specifically, at 2 or 3 dpi of the ΔCRISPR strain we typically observed an increase in phage densities. Analysis of the infectivity patterns of these phages on the different host backgrounds (WT, ΔCRISPR) revealed that this increased density was due to the emergence of escape phages that had evolved to overcome MADS (Fig. 4g, red circles). Evolution of these escape phages during infection of the ΔCRISPR strain was identified in 20 out of 42 (47%) independent experiments spanning different initial phage inoculi, from 10^4^ PFU/mL to 10^9^ PFU/mL (corresponding to MOI of 10^-3^ to 10^2^). By contrast, when bacteria carried both the MADS and the CRISPR-Cas systems (i.e., SMC4386-WT), analysis of the infectivity patterns of the phage populations on the different host backgrounds (WT, ΔCRISPR) detected escape phages in 12% of the replicates (5 out of 42) and only in experiments where the initial phage titres were very high, that is 10^9^ PFU/mL (MOI 10^2^, Fig. 4h, red circles). However, recovery of phage population density was only observed in 1 out of these 5 replicates (Fig. 4h). Collectively, these experiments support the model prediction that evolution of Acr-phages to overcome MADS is suppressed when bacteria also encode a CRISPR-Cas immune system, since imperfect infectivity of Acr-phage on CRISPR-immune bacteria limits both the emergence and spread of such epigenetic mutants of the phage.

## Discussion

For the past few years, new defence systems have been discovered at unprecedented rates, suggesting that many more remain to be uncovered. These discoveries have mainly been based on computational “guilt-by-association” approaches which take advantage of the fact that defence genes often cluster together in bacterial and archaeal genomes in so-called “defence islands”^14,17,23^. Anti-phage genes are also frequently identified in MGE, where they form hotspots of genetic variation within the MGE^37–39^. In addition, these MGE tend to integrate at preferential locations in bacterial genomes, known as integration hotspots. Therefore, systematic mapping of integration hotspots and careful analyses of their gene content is becoming a valuable strategy for the discovery of the full repertoire of bacterial defence systems^40–43^. We now understand that bacteria encode many more defence systems than previously thought, and that individual bacterial genomes typically contain multiple defences. However, how these defences interact at the molecular, transcriptional and phenotypic levels, and how this shapes bacteria-phage coevolution remains largely unclear.

Here, we identify a new defence system that we named MADS in a genomic locus that is a hotspot for defence systems. MADS uses N6-methyladenosine modification of a specific recognition sequence to discriminate self from non-self DNA. We found that a bacterial strain that encodes MADS alone suppresses phage DMS3*vir* densities but only transiently as phages can readily acquire epigenetic modifications to escape immunity mediated by MADS. Interestingly, when bacteria also carried CRISPR-Cas immunity, phage escape evolution was rarely observed, despite the phage carrying an anti-CRISPR gene that blocks the CRISPR-Cas immune system of the host. Mathematical modelling explains why this synergy between the two defence systems emerges. It has previously been shown that phage carrying *acr* genes need to cooperate in order to overcome CRISPR immunity of their host bacteria, and this in turn causes amplification only if the initial phage densities are sufficiently high^10,11^. Cooperation is needed because initial infections of CRISPR immune bacteria by Acr-phages often fail, due to phage genomes being detected and destroyed by CRISPR immune complexes before they are inactivated by the Acr proteins. However, some Acr proteins will have been produced, and although the initial infection was unsuccessful, the cell enters an immunosuppressed state that can be exploited in a secondary infection (Extended Data Fig. 6). The probability that this cell will be re-infected by another Acr-phage increases with phage density, explaining the critical tipping point in Acr-phage density beyond which phage amplification is observed. However, in the presence of a second layer of defence, such as MADS, these cells with a suppressed CRISPR-Cas immune system are unavailable for phage replication. To exploit these cells, phages first need to acquire the epigenetic modification to overcome MADS. The presence of CRISPR-Cas immunity limits the evolution of Acr-phage mutants that escape MADS, and even if phage mutants arise, they cannot spread as efficiently as they would in the absence of CRISPR-Cas immunity (Extended Data Fig. 6). Based on our understanding of the synergy between CRISPR-Cas and MADS upon infection with Acr-phages, we expect that similar synergistic interactions may occur between CRISPR-Cas and other innate immune systems. Future work will be needed to test this hypothesis, and if validated, to understand if and how the activity of these systems is coordinated to provide an optimal anti-phage response.

Collectively, our study sheds light on how multi-layered defence systems can shape bacteria-phage interactions, and shows that the coexistence of CRISPR-Cas and MADS impairs the successful deployment of phage counter-defence strategies by interfering with phage cooperation. Specifically, the presence of MADS reduces the infection success of immunosuppressed bacteria by Acr-phages, whereas the presence of the CRISPR immune system limits the emergence and spread of epigenetic mutants of the phage that overcome MADS. A more detailed understanding of the interactions between multi-layered defences and the counter-defences encoded by phages will be key to predict and manipulate bacteria-phage interactions in natural and clinical settings.

## Supporting information

PADLOC_output_SMC4386

Island_boundaries_genomes

DF_output_defence_islands

Primers_list

## Acknowledgments

A.M. was supported by funding from a PhD studentship sponsored by the College of Life and Environmental Sciences (CLES, University of Exeter), and the Centre for Environment, Fisheries and Aquaculture Science (CEFAS). This work was supported by a grant from the ERC (ERC-STG-2016-714478 - EVOIMMECH), which was awarded to E.R.W. A.C. received funding from the European Union’s Horizon 2020 research and innovation programme under the Marie Skłodowska-Curie grant agreement No 834052 and from IdEx Université Paris Cité (ANR 18 IDEX 0001). K. B. and C.C. drew on funding from the MRC MR/R020787/1. S.v.H. acknowledges funding from the Biotechnology and Biological Sciences Research Council (BB/S017674/1). The authors thank Dr. L. Zhang for his support in optimizing the protocol for arbitrary PCR, Dr. D. Walker-Sünderhauf for assistance in generating the transposon-mutant library, Dr. B. Watson for contributing to phage DNA isolation, Dr. A. Bernheim for feedback on building macromolecular models to detect MADS, E. Cenraud for DefenseFinder analyses, Prof. Bondy-Denomy for kindly providing phage DMS3-Gm, Prof. R. Lavigne and Dr. J. Wagemans for kindly providing phages LPB1, LMA2 and LUZ24, Prof. O’Toole for kindly providing phage DMS3*vir* and strain *P. aeruginosa* UCBPP-PA14 *csy3::lacZ*, and Prof. Davidson for kindly providing strain SMC4386.

## Authors contributions

Conceptualization, A.M., A.C. and E.R.W.; Methodology, A.M., E.P., C.C., S.G, K.B., A.C., and E.R.W.; Investigation A.M., E.P., C.C., S.G, R.C., B.J.P., M.C., A.G., S.P., K.B., S.v.H., A.C. and E.R.W.; Formal Analysis, A.M., E.P., C.C., S.G, S.P., K.B., A.C., and E.R.W.; Mathematical Modelling, S.G; Writing – Original Draft, A.M., A.C. and E.R.W.; Writing – Review & Editing, A.M., A.C. and E.R.W. with contributions from E.P., C.C., S.G., R.C., B.J.P., M.C., S.P., K.B., S.v.H.; Supervision, A.C. and E.R.W.; Funding Acquisition, E.R.W.

## Methods

All experiments were carried out in six biological replicates unless otherwise specified.

### Bacterial strains

*P. aeruginosa* UCBPP-PA14 *csy3::lacZ* (referred to as PA14ΔCRISPR, since it carries a disruption of an essential *cas* gene that causes the CRISPR-Cas system to be non-functional), was grown overnight at 28°C or 37°C in LB broth. *P. aeruginosa* SMC4386 (referred to as SMC4386-WT), and deletion mutants of this strain (referred to as ΔCRISPR, ΔMADS, ΔCRISPRΔMADS, Δ*mad1*, Δ*madi2*, Δ*mad3*, Δ*mad4*, Δ*mad6*, Δ*mad7,* Δ*mad8*) were growth at 28°C or 37°C in either LB broth or M9 medium (22 mM Na_2_HPO_4_; 22 mM KH_2_PO_4_; 8.6 mM NaCl; 20 mM NH_4_Cl; 1 mM MgSO_4_; 0.1 mM CaCl_2_) supplemented with 0.2% glucose. *E. coli* MFD*pir* was used as donor to build the transposon mutant library, *E. coli* DH5α was used to assemble the allelic exchange vectors for the gene deletions, and *E. coli* S17.1λ*pir* was used as donor strain to deliver the allelic exchange vectors to recipient cells during conjugation assays. Whenever applicable, media was supplemented with ampicillin (100 µg/mL), streptomycin (50 µg/mL), or gentamicin (either 30 µg/mL, when selecting for *E.coli*, or 50 µg/mL, when selecting for *P. aeruginosa*) to ensure plasmid maintenance. *E. coli* MFD*pir* was cultured in the presence of diaminopimelic acid (DAP) (0.3 mM).

### Phage strains

Recombinant temperate phage DMS3-Gm, which encodes a gentamycin resistance gene, was used to enable selection of lysogens following infection of WT and mutant SMC4386 strains as well as the transposon mutant library. Phage DMS3*vir*, which is obligately lytic due to the deletion of the c-repressor gene, and/or phage DMS3*vir*Δ*acrIE3*, which lacks both the c-repressor gene and the anti-CRISPR gene that blocks the Type IE CRISPR-Cas system of strain SMC4386, were used in all other experiments, and have been described previously^28,44^. Lytic phages LMA2, LUZ24 and temperate phage LPB1 were used in spot assays to determine the range of resistance conferred by MADS. Phage stocks were extracted from lysates prepared on *P. aeruginosa* PA14ΔCRISPR or SMC4386ΔCRISPR and stored at 4°C.

### Adsorption assay

The measurement of phage adsorption was performed according to Kropinsky, 2009^45^ with some modifications. Briefly, 9 mL of *P. aeruginosa* SMC4386-WT and PA14ΔCRISPR culture (OD_600_=0.25) were infected with 2×10^6^ plaque forming units (PFU)/mL of phage DMS3*vir*. Every 6 min for 1h, an aliquot of 100 µL was taken from each vial and transferred into a chilled Eppendorf tube containing 800 µL of LB broth and 100 µL of chloroform. Extracted phages were serially diluted and spotted onto a lawn of *P. aeruginosa* PA14ΔCRISPR to determine the phage titre. The experiment was carried out in a water bath, at 28°C and with 100 rounds per minute (rpm) agitation.

### Efficiency of Plaquing (EOP) assays

EOP assays were carried out on square polystyrene plates containing LB with 1.5% agar. A mixture of molten soft LB agar (0.5%) and 300 µL of bacteria (grown overnight in LB broth) were poured on top of the hard agar layer and allowed to set. Next, 5 µL of serially diluted phage were spotted on the resulting plates, which were subsequently incubated overnight at 28°C and plaque forming units (PFUs) were enumerated the next day. The EOP was determined as the ratio of the number of PFUs on a mutant *P. aeruginosa* SMC4386 and the *P. aeruginosa* SMC4386-WT strains. To be able to calculate the EOP with DMS3*virΔacr*, we arbitrarily set the phage titre on *P. aeruginosa* SMC4386-WT at 1 PFU/mL (instead of 0).

### Generation of the transposon (Tn)-mutant library

To generate the transposon-mutant library we used the synthetic construct pBAM (born-again-mini-transposon) described by García et al., 2011^46^. Specifically, we utilised the vector pBAMD1-4^47^, which delivers the Tn5 while conferring resistance to streptomycin to the target cells at the same time, allowing for selection of transconjugants. The pBAMD1-4 vector was delivered to the recipient bacteria via conjugation using the *E. coli* MFD*pir* strain as donor cell, following previously described methods^48^. Briefly, we separately incubated 15 mL of *E. coli* MFD*pir* donor cells and recipients (*P. aeruginosa* SMC4386-WT and SMC4386ΔCRISPR strains), which were incubated overnight at 37°C with agitation at 180 rpm. Recipients were grown in LB broth, while donors were grown in LB broth supplemented with DAP (0.3mM), ampicillin (100 µg/mL) and streptomycin (50 µg/mL). The following day, 10 mL of each strain was pelleted and washed twice with 10 mL of M9 salts saline solution. Bacterial cultures where pelleted again and resuspended in 10 mL of LB supplemented with 0.3 mM DAP. Donors and recipient were mixed together (7500 µL donor: 500 µL recipient), pelleted and resuspended in 1 mL of M9 saline solution. To allow for conjugation between donor and recipient, 100 µL of mix was spotted onto 10 individual Binder-Free Glass Microfiber 1.2 µM filter papers (Whatman) which were placed onto a squared LB agar plate supplemented with DAP (0.3mM), and incubated at 28°C for 48h. Cells from each filter were then recovered into LB supplemented with streptomycin (to kill the non-transconjugants), pelleted, washed twice (resuspended first in 1 mL and then in 100 µL) and plated onto an LB agar plate supplemented with streptomycin to select for the transconjugants. Plates were incubated for 24-48h at 28°C. For each recipient strain, an *E. coli* MFD*pir* donor strain without plasmid was used as negative control. Based on pilot data that yielded around 1000 transconjugants, this procedure was carried out in 10 independent biological replicates, which were pooled during the last step in order to obtain a saturated Tn mutant library.

### Measuring frequencies of lysogeny

#### Lysogeny in PA14ΔCRISPR, SMC4386 and derivative knockouts strains

Cultures were grown overnight at 37°C with 180 rpm agitation, in 6 mL of LB broth. Cultures were then diluted 1:100 into fresh LB broth and infected with phage DMS3-Gm^49^ at an MOI of 0.01. After 24h, samples were serially diluted, plated onto selective (LBA with Gm) and non-selective (LBA) agar plates and incubated overnight at 28°C. Colonies were enumerated the next day. The proportion of lysogens was expressed as the number of CFU (Colony Forming Units) grown on selective plates divided by the number of CFU on non-selective media.

#### Lysogeny in the Tn5 mutant library

The transconjugants (see **Generation of the transposon (Tn)-mutant library** for details on how they were obtained) were scraped off LBA plates, pooled, and resuspended in 10 mL of LB broth and then infected with 10^5^ PFU of phage DMS3-Gm, followed by overnight incubation at 28°C with agitation at 180 rpm. Each day and for 3 days, 1 mL of the culture was transferred into 10 mL of fresh LB broth, the phages were extracted to monitor phage titre and bacteria were plated onto non-selective LBA as well as LBA supplemented with streptomycin (selecting for Tn mutants) or both streptomycin and gentamicin (selecting for Tn mutants carrying the DMS3-Gm prophage). Plates were incubated at 28°C for 24-48h. Lysogenization of bacterial colonies on LBA with streptomycin and gentamicin was confirmed by colony PCR using primers Crep_F (forward, 5’-GCGGAATGAGCGCTAAACC-3’) and Crep_R (reverse, 5’-CAAGTGCTTTAGCGAGGAATGC-3’), that amplify the c-repressor gene of phage DMS3.

### Localization of the Tn5 insertions

Before being subjected to arbitrary PCR (described below), we verified using colony PCR that the clones of interest carried the miniTn5 insertion using the primer pairs PS5 (5’-CCCTGCTTCGGGGTCATT-3’) and PS4 (5’-CCAGCCTCGCAGAGCAGG-3’), and PS5 and PS6 (5’-GGACAAATCCGCCGCCCT-3’), using cells carrying the plasmid pBAMD1-4 as a positive control. Both primer pairs amplify the *oriT* region of the plasmid, leading to a product size of 225 bp and 665 bp respectively. Only in case of absence of *oriT* amplification we proceeded with the arbitrary PCR. Next, we applied a protocol of arbitrary PCR to identify the location of Tn5 insertions, based on the methods described by García et al., 2014^47^ and Saavedra et al., 2017^50^. In the first round of arbitrary PCR we used the forward primer ME-O-Sm-Ext-F (5’-CTTGGCCTCGCGCGCAGATCAG-3’) and the reverse primer ARB6 (5’-GGCACGCGTCGACTAGTACNNNNNNNNNNACGCC-3’), while in the second round of arbitrary PCR we used the forward primer ME-O-Sm-Int-F (5’-CACCAAGGTAGTCGGCAAAT-3’) and the reverse primer ARB2 (5’-GGCACGCGTCGACTAGTAC-3’). We followed the PCR conditions that have been previously described^47^. Since all the transconjugants are different from one another (i.e., the Tn5 is in different location in the bacterial genome), some PCR conditions, such as the annealing temperature, work for some clones but not for others. Therefore, for clones where we did not obtain clear and definite bands after the second round, we adjusted the protocol as follows: in the first round the number of cycles increased from 30 to 35 and/or the annealing temperature gradually increased to a maximum of 38°C; in the second round the annealing temperature increased to 52°C. PCR products obtained after the second round of arbitrary PCR were gel purified and sent for Sanger sequencing (Eurofins Genomics UK Limited, Wolverhampton, UK). The chromatogram derived from the Sanger sequencing was mapped against the genome of *P. aeruginosa* SMC4386-WT using Geneious v10.2.6 to identify the genes where the transposon was inserted.

### Generation of gene knockouts

The deletion of CRISPR-Cas and MADS systems, as well as the deletion of single genes from MADS were carried out using two-step allelic exchange, as described by Hmelo et al., 2015^51^. The homologous sequences flanking either side of the desired target system and/or gene were synthesized by Integrated DNA Technology (IDT™) or PCR amplified and fused together via SOE-PCR^52,53^, and then cloned into the pDONRPEX18Gm donor vector via Gateway cloning. The resulting allelic exchange vector was transformed into chemically competent *E. coli* DH5α and verified by PCR. Vectors were then electroporated into competent *E.coli* S17-1λ*pir* to allow for conjugation with *P. aeruginosa* SMC4386-WT recipient strains^51^. The merodiploids that were obtained were selected on cetrimide agar plates (to select for *P. aeruginosa*) supplemented with gentamicin (50 µg/mL). Every genomic deletion was confirmed by colony PCR, first by positive amplification of the knockout junction, and then by negative amplification of the left and right side of the intact system and/or gene, followed by Sanger sequencing (Eurofins Genomics UK Limited, Wolverhampton, UK). The list of primers used to generate and screen the knockout strains are listed in Supplementary Table 4.

### Bioinformatic analysis of the distribution of MADS

#### Analysis of protein domains and HMM profiles

MADS locus protein sequences were determined from genome annotations made using Prokka v1.14.6, which uses Prodigal to predict protein regions. These proteins were searched against the pfam (protein family) database, as well as Phyre2 and HHpred, to perform HMM-HMM matching-based remote homology detection and obtain structural predictions^54,55^. Curated alignments for each gene were constructed by identifying homologues for each protein sequence using HHpred. A probability cut-off score of 50% was set, with default parameters for all other options, searching against the PDB database. Alignments extracted from HHpred outputs (max 250 sequences) were then used to generate profile-HMMs using the *hmmbuild* function from HMMER v3.0. For further investigations of remote homology, HHpred was run against the PDB, COG, NCBI conserved domains and Pfam-A databases.

#### Genomes

A total of 172,366 bacterial and archaeal RefSeq assemblies (retrieved January 2022 using ncbi-genome-download, https://github.com/kblin/ncbi-genome-download/) were downloaded for use in prevalence analysis.

#### Macromolecular models for MADS

MacSyFinder v2, a tool for macromolecular systems detection^56^, was used to develop models for MADS. This tool requires the specification of mandatory or accessory components within the system, which are not biological definitions but rather describe whether a protein is easy to detect or more divergent and thus harder to detect with a single HMM profile, as well as whether components are frequently missing from systems.

After making sure that no known anti-phage systems could be identified by PADLOC^57^ and DefenseFinder^58^ within the MADS operon (Supplementary Table 1), genomes were retrieved in genbank (.gbff) format and ordered protein fasta files were created by extracting the CDS features using SeqIO from Biopython. Initial models with various levels of system completeness and genomic distance between components were tested. Initially, all *P. aeruginosa* RefSeq genomes (n=6,103) were searched. Next, preliminary tests, and manual inspection of hits from a subset of the entire Bacterial and Archaeal RefSeq dataset (n=15,000), the following model parameters were set. Proteins MAD6, 7 and 8 were defined as mandatory and their presence was required for the system to be detected. Proteins MAD1, 2 and 5 were defined as accessory due to their homology to Restriction-Modification system components and widespread regulators, which could lead to false positive hits if made mandatory. Finally, proteins MAD3 and 4 were also classified as accessory, due to them not being reliably detected in all systems. In addition, the maximum inter gene distance was set to 10.

#### Taxonomic distribution and plots

Species classifications for genomes with MADS were retrieved using the Entrez python module. Taxonomic trees were retrieved from NCBI using the ete3 v3.1.2 module, and used to visualise the phylogenetic distribution of MADS. Other plots were created in R, with operons plotted using gggenes. Additional editing of plots was performed using Inkscape v1.2.

#### Reproducibility and computational resources

The University of Exeter’s Advanced Research Computing Facilities were used to carry out for this bioinformatics analysis. A Snakemake pipeline was used to run MacSyFinder searches on all genomes, whilst the remaining analyses were carried out using R v4.0.4 and Python version 3.9.7 with Biopython v1.79. All scripts used for this analysis are available at https://github.com/elliekpursey/Maestri.

### Bioinformatic analysis of defence islands

A total of 454 publicly available complete *P. aeruginosa* genomes were included in the assessment of the putative defence island (Supplementary Table 2). Defence islands were extracted from genomes by first examining the BLASTx percentage identity and genome coordinates against relevant query reference sequences of the distinct gene boundaries pheT (MPAO1_RS11510) and a histidine kinase (MPAO1_RS11445)^59^. FASTA sequence files of defence islands were extracted from complete genome FASTA files using BEDTools v2.29.2 getfasta and subtract, based upon BLASTx genome coordinates^60^. Prokka v1.14.6 was used to annotate defence islands and the remainder of the genome^61^. Defence systems in all sequence files were identified using using PADLOC^57^ and DefenseFinder^58^, both with additional CRISPR array detection. Wilcoxon’s signed rank testing was used to determine enrichment for defence systems on the island and was performed using R v4.1.3. Manual inspection of the defence hotspots containing MADS was carried out using Geneious v10.2.6

### Growth curves

Growth curves to assess the role of MADS during infection with phage DMS3*vir* were performed in 96-well plates with agitation at 37°C. Briefly, overnight cultures of WT and mutant strains of *P. aeruginosa* SMC4386 grown in LB media at 37°C with agitation at 180 rpm were diluted 100-fold into fresh LB media and grown until cultures reached mid-log phase (OD600 nm of 0.3, approximately 10^8^ CFU/mL). One mL of each mid-log culture was centrifuged for 3 min at 6000 rpm to remove the supernatant and the pellets were infected with 1 mL of DMS3*vir* at an MOI of 10 in LB and incubated for 10 min at 37°C whilst shaking at 180 rpm. After initial incubation to synchronise the infection, 200 µL of the infected cultures were transferred to a 96-well plate, then 20 µL of mineral oil were added on the surface of each well to avoid evaporation and bacterial growth was measured by optical density at 600 nm (OD600) for 24h at 37°C with agitation in a BioTek Synergy 2 plate reader. At t=0 200 µL samples were used to quantify initial bacteria CFU/mL and phage titre PFU/mL. Growth measurements in the absence of phage were carried out in parallel as a control.

### Analysis of MADS self/non-self discrimination

#### Infection assays in liquid medium

Infection assays to measure the population dynamics of phage DMS3*vir* in the presence or absence of MADS were performed in glass vials by inoculating 6 mL M9 medium supplemented with 0.2% glucose with approximately 5 x 10^7^ CFU bacteria from fresh overnight cultures (also grown in M9 medium + 0.2% glucose) of either *P. aeruginosa* SMC4386-WT strain or the isogenic ΔCRISPR, ΔMADS or ΔCRISPRΔMADS strains. Cultures were infected at MOI 0.1 and 10, incubated at 37°C while shaking at 180 rpm and transferred daily (1:100 dilution) for three days into fresh medium. Phages were chloroform extracted (1:10 volume) every day and their titres measured via spot test assay onto a lawn of the sensitive *P. aeruginosa* PA14ΔCRISPR strain. Plates were incubated overnight at 28°C. We monitored whether phages in these experiments evolved to overcome bacterial defence systems, and whether this was due to genetic or epigenetic mutations. To this end, we performed plaque-purification of phages from the SMC4386 lawns using chloroform extractions, and phage were titrated on lawns of SMC4386-WT and PA14ΔCRISPR strains. These phages were then used in a next round of parallel infections on SMC4386-WT or PA14ΔCRISPR strains as hosts, at 28°C in glass vials containing 6mL of LB broth, while shaking at 180 rpm. Phages were chloroform extracted again, and titrated on both strains. This process was repeated for another round of infection, to understand the heritability of the escape phenotype of the phage.

#### Phage and bacteria DNA extraction for sequencing

For phage DNA extraction the 4 replicate experiments of DMS3*vir* that during infection assays with *P. aeruginosa* SMC4386ΔCRISPR (referred to as ΔCRISPR) gained the capability to amplify after 2 days of infection (see Fig.3a), were amplified on ΔCRISPR and ΔCRISPRΔ*mad2*, which is the mutant lacking the predicted methylase, while the ancestral DMS3*vir* was amplified on ΔCRISPRΔMADS. To this end, 500 μL of bacteria from a fresh overnight culture were inoculated into 50 mL of LB broth and mixed with 100 μL of an approximately 1×10^8^ PFU phage stock. Those infected cultures were grown overnight in 50 mL of broth at 28°C, 180 rpm. The resulting viscous cultures were centrifuged at 25,000 x*g* and phages were found to have concentrated in the pellet. The pellet was resuspended with 5 mL of phosphate buffered saline (PBS) and centrifuged at 21,000 x*g* to remove bacterial cells. Phages were concentrated from the supernatant using 0.5 mL 100kDa Amicon spin filters as per the manufacturers guidelines and retained in the filters. Whilst still in the filter, phages were washed twice with DNase I buffer (up to 400 µL), treated with DNase I as per the manufacturers guidelines, washed twice with RNase buffer (up to 400 µL), treated with RNase A as per the manufacturers guidelines, washed twice with RNase buffer (up to 400 µL) and eluted from the Amicon filters by inversion and centrifugation with RNase buffer (up to 400 µL) to give a final volume for each phage solution of 400 µL. Phage DNA was extracted using a proteinase K lysis step (20 µg of proteinase K, 0.5% SDS, 20mM EDTA pH 8.0) for 1 h at 60 °C followed by 2 phenol:chloroform extractions (1:1 volume, inversion and centrifugation), DNA was precipitated using sodium acetate (1/10 volume, 3M, pH 7.5 and ethanol (2.5 volume, 100%, ice-cold) overnight at -20°C. DNA was pelleted (30min, 20,000 x*g*, 4°C) and washed twice with ice-cold 70% ethanol, dried and resuspended in TE buffer.

For bacterial DNA extraction (*P. aeruginosa* SMC4386-WT, SMC4386ΔCRISPR, SMC4386ΔCRISPRΔMADS, SMC4386ΔCRISPRΔ*mad2*) bacteria were pelleted from overnight culture and DNA was extracted using the phenol chloroform method outlined previously. Quality Control and quantification of bacteria and phage DNA was performed with NanoDrop, Qubit and agarose gel electrophoresis; 2 µg of DNA were used for Pacific Biosciences (PacBio) sequencing (Centre For Genomic Research, Liverpool, UK).

#### PacBio sequencing

Barcoded SMRT-Bell PacBio libraries were created from each DNA sample and run on a SMRT cell in CLR mode on a Sequel IIe platform, yielding >300,000x coverage for phage samples and 1,400 – 2,200x coverage for bacterial samples. The PacBio SMRT-Link v10.0 analysis pipeline (https://www.pacb.com/support/software-downloads/) was used to demultiplex samples, to detect methylation signals based on polymerase kinetics, and to identify motifs associated with methylation, all under default parameters. Resulting diagnostic plots and gff files were inspected manually. Output files, including logs, are available at https://github.com/scottishwormboy/Maestri_pbio. The genome reference used for DMS3 was NC_008717.1. For SMC4386, the PacBio reads were used to create a new genome reference (accession: PRJNA905210) from the wild-type, using SMRT-Link v10.0 for microbial assembly.

### CRISPR immunosuppression assay

CRISPR immunosuppression assays were performed as previously described^36,53^. This assay relies on transformation of *P. aeruginosa* SMC4386 cells with plasmid pHERD30T (non-targeted by the SMC4386 CRISPR-Cas system) and pHERD30T-cr2sp1-SMC (targeted by SMC4386 CRISPR-Cas system). Briefly, the pHERD30T-cr2sp1-SMC plasmid was constructed by inserting a 32 nucleotide protospacer matching the 1^st^ spacer of CRISPR array 2 of the *P. aeruginosa* SMC4386 strain, flanked by the AAG Protospacer Adjacent Motif (PAM). Oligonucleotides containing the PAM and protospacer sequence (5’-agctt<u>AAG</u>AACCTCTACGAGCAGACCGAGTTGAAAGGGCAg-3’ and 5’-aattcTGCCCTTTCAACTCGGTCTGCTCGTAGAGGTT<u>CTT</u>a-3’, restriction sites overhangs are indicated in small caps, protospacer in capitals and PAM underlined) were annealed to create overhangs compatible with HindIII and EcoRI, phosphorylated by T4 Polynucleotide Kinase and ligated in EcoRI-HindIII digested pHERD30T vector. Cultures of *P. aeruginosa* SMC4386-WT and isogenic ΔCRISPR, ΔMADS and ΔCRISPRΔMADS strains grown overnight in 50 mL of LB medium (approximately 3.5 x 10^9^ CFU/mL) were divided in three 50 mL tubes with 10 mL of culture in each. Cells were either non-infected or infected using a final density of 10^9^ PFU/mL (MOI=0.3) of DMS3*vir* or DMS3*vir*Δ*acrIE3*. After 2h of incubation at 37°C with agitation at 180 rpm, cells were harvested by centrifugation at 3500 rpm for 15 minutes. A sample of the supernatant was kept for phage titration by spot assay to validate homogeneity of the phage titre across the replicates. Cells were made electrocompetent by washing them twice with 1 mL of 300 mM sucrose solution at room temperature and resuspended in 300 µL of the same solution. A 100 µL sample of the resuspended cells was taken to evaluate the bacterial density. The remaining 200 µL were equally divided across two vials and electroporated with 500 ng of either pHERD30T or pHERD30T-cr2sp1-SMC, followed by addition of 700 µL fresh LB medium. After incubating for 1h at 37°C at 180 rpm, bacteria were pelleted, resuspended in 100 µL of LB medium and serially diluted in LB-medium. Fifty microliters of each dilution were spotted on LB agar plates containing Gentamycin (50 µg/mL) and incubated overnight at 37°C to allow transformants to grow. Relative transformation efficiency was calculated as the number of colonies obtained after transformation with pHERD30T-cr2sp1-SMC divided by the number of colonies obtained after transformation with pHERD30T.

### Plasmid transformation assay

This assay was adapted from the CRISPR immunosuppression assay to fit the MADS. It relies on transformation of *P. aeruginosa* SMC4386 cells with plasmid pHERD30T (non-targeted by the SMC4386 MADS) and pHERD30T-M24 (targeted by SMC4386 MADS). Briefly, the pHERD30T-M24 plasmid was constructed by inserting a 11 nucleotide MADS target sequence (5’-TCAGCGCGTCC-3’). Oligonucleotides containing the target sequence (5’-agcttTCAGCGCGTCCg-3’ and 5’-aattcGGACGCGCTGAa-3’, restriction sites overhangs are indicated in small caps and target sequence in capitals) were annealed to create overhangs compatible with HindIII and EcoRI, phosphorylated by T4 Polynucleotide Kinase and ligated in EcoRI-HindIII digested pHERD30T vector. Each of the two plasmids were then transformed in either *P. aeruginosa* SMC4386ΔCRISPRΔMADS or isogenic ΔCRISPRΔ*mad3*. These transformed strains were then used for plasmid production, which were purified with GeneJET Plasmid Midiprep kit (Thermo Scientific, USA) according to the manufacturer’s instructions. Plasmid produced in *P. aeruginosa* SMC4386ΔCRISPRΔMADSwere called pHERD30T-Nonmeth and pHERD30T-M24-Nonmeth, while plasmids produced in *P. aeruginosa* SMC4386ΔCRISPRΔ*mad3* were called pHERD30T-Meth and pHERD30T-M24-Meth.

Cultures of *P. aeruginosa* SMC4386-WT and isogenic ΔCRISPR, ΔMADS and ΔCRISPRΔMADS strains grown overnight in 20 mL of LB medium (approximately 3.5 x 10^9^ CFU/mL) were harvested by centrifugation at 3500 rpm for 15 minutes. Cells were made electrocompetent by washing them twice with 2 mL of 300 mM sucrose solution at room temperature and resuspended in 500 µL of the same solution. A 100 µL sample of the resuspended cells was taken to evaluate the bacterial density. The remaining 400 µL were equally divided across four vials and electroporated with 500 ng of either pHERD30T-Nonmeth, pHERD30T-M24-Nonmeth, pHERD30T-Meth or pHERD30T-M24-Meth, followed by addition of 700 µL fresh LB medium. After incubating for 1h at 37°C at 180 rpm, bacteria were pelleted, resuspended in 100 µL of LB medium and serially diluted in LB-medium. Fifty microliters of each dilution were spotted on LB agar plates containing Gentamycin (50 µg/mL) and incubated overnight at 37°C to allow transformants to grow. Relative transformation efficiency of methylated or non-methylated plasmids (respectively) were calculated as the number of colonies obtained after transformation with pHERD30T-M24-Nonmeth or pHERD30T-M24-Meth divided by the number of colonies obtained after transformation with pHERD30T-Nonmeth or pHERD30T-Meth, respectively.

### Phenotypic interactions between CRISPR-Cas and MADS

#### Tipping-point assay

Infection assays were performed in glass vials by inoculating 6 mL M9 medium supplemented with 0.2% glucose with approximately 10^7^ CFU bacteria from fresh overnight cultures (also grown in M9 medium + 0.2% glucose) of the *P. aeruginosa* SMC4386-WT strain and the isogenic ΔCRISPR, ΔMADS and ΔCRISPRΔMADS strains. To these vials, phage DMS3*vir* was added at a MOI of 10^-4^, 10^-3^, 10^-2^, 10^-1^, 1, 10^1^, or 10^2^, and incubated at 37°C while shaking at 180 rpm. Cultures were transferred daily (1:100 dilution) for three days into fresh M9 medium. For the *P. aeruginosa* SMC4386ΔMADS and the SMC4386ΔCRISPRΔMADS strains we performed additional infections at MOI 10^-5^ and MOI 10^-6^ for seven days. Phages were chloroform extracted every day and the titres were measured using spot assays onto lawns of each of the four different bacterial strains (i.e., *P. aeruginosa* SMC4386-WT, ΔCRISPR, ΔMADS and ΔCRISPRΔMADS strains). Plates were incubated overnight at 28°C.

To identify replicate experiments in which phage mutants had emerged that escape MADS, we determined for each phage sample the EOP ratio’s on *P. aeruginosa* SMC4386ΔCRISPRΔMADS and either the ΔCRISPR or WT bacteria. For WT DMS3*vir* phages, these ratios are 127 ±17 and 142 ± 33 (mean ± 95% c.i.), respectively. For clonal phage populations of DMS3*vir* escape mutants (carrying the epigenetic modification to overcome MADS), these ratios are 0,91 ± 0,15 and 1,18 ± 0,12 (mean ± 95% c.i.), respectively. We therefore applied a conservative threshold of EOP=5 (i.e. corresponding to ± 20% escape mutants in the phage population) to identify samples that contained significant numbers of MADS escape phages. Replicates where escape phages were detected are depicted as red circles in Figure 4 g, h.

### Mathematical modelling

#### Epidemiological dynamics (no phage evolution)

We develop a model to understand the dynamics of bacteriophages in a multiresistant bacteria population. Earlier studies have examined the evolution of Acr in well-mixed environments^44,62^. Here, we explore how the addition of a second resistance could affect the dynamics.

Extended Data Fig.5a shows a schematic representation of the phage life cycle (where we assume that bacteria are initially CRISPR resistant to the phage). The Acr-phage is able to infect resistant bacteria with a probability 1 − *ρ*, where *ρ* is a measure of CRISPR efficiency. These infections can either lead in (i) the production of immunosuppressed cells with probability *r*(1 − *ϕ*), where *r* is is another measure of CRISPR efficiency and *ϕ* is a measure of anti-CRISPR efficiency, or (ii) they can result in cell lysis and the release of *B* virions with probability =1 − *r*(1 − *ϕ*)>(1 − *m*), where *m* is a measure of MADS efficiency.

This yields the following system of ordinary differential equations (where *N* = *R* + *S*):

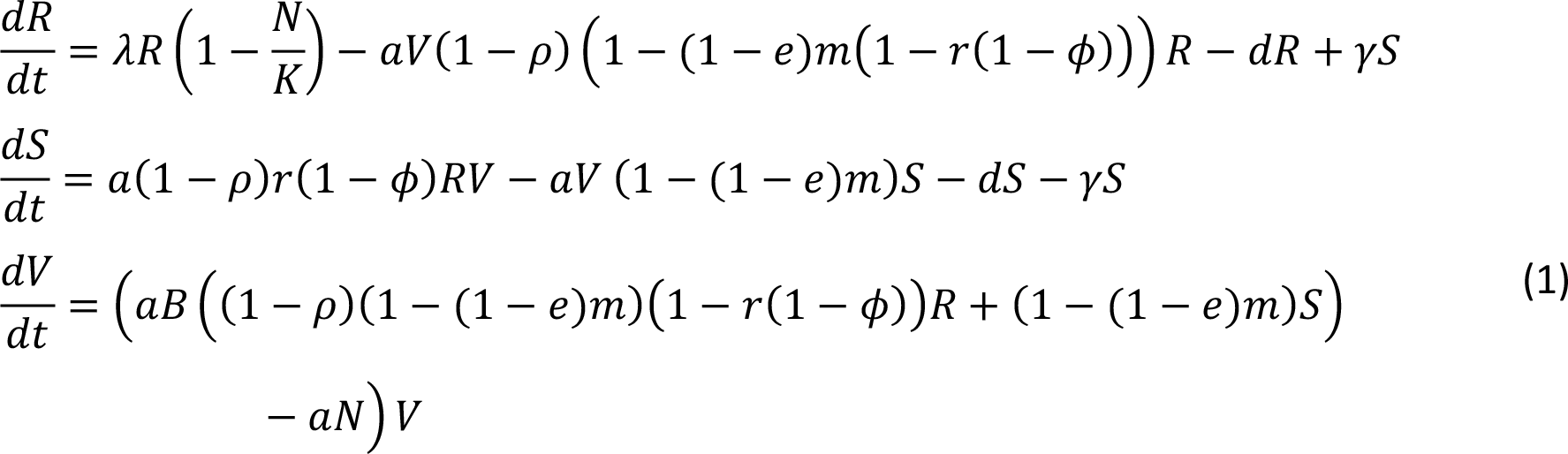

The change in the total density of virus *V* at the beginning of an epidemic where *S* = 0 and *N* = *R*:

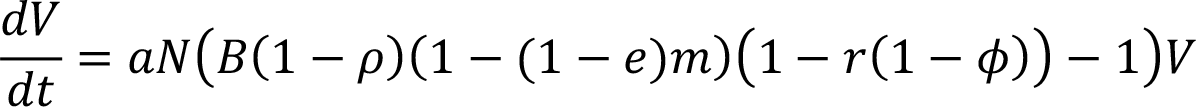

In other words, the phage population can grow only when:

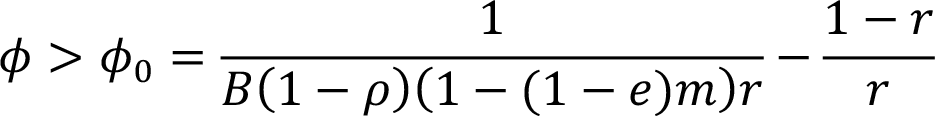

Yet, if one introduces a large density of phages in the host population they will immunosuppress a fraction *S*⁄*N* of the cells. This will yield the following threshold

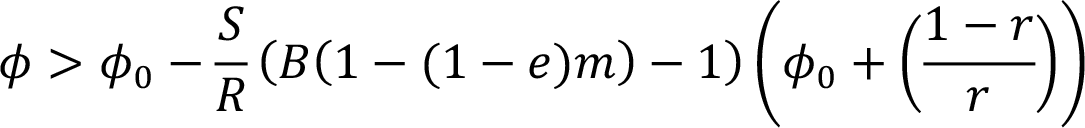

In other words, we recover the results of ref^62^ (in the case where *m* = 0) and extend it to the case where bacteria carry the MADS resistance. The above expression shows that increasing *m* always increases the threshold density of viruses above which the epidemic can take off (see Fig. 4a).

#### Evolutionary dynamics of the phage (MADS escape)

In the following we consider an alternative model where the phage can acquire an epigenetic mutation allowing the virus to escape MADS (i.e., the parameter *e* = 1 for the escape mutant):

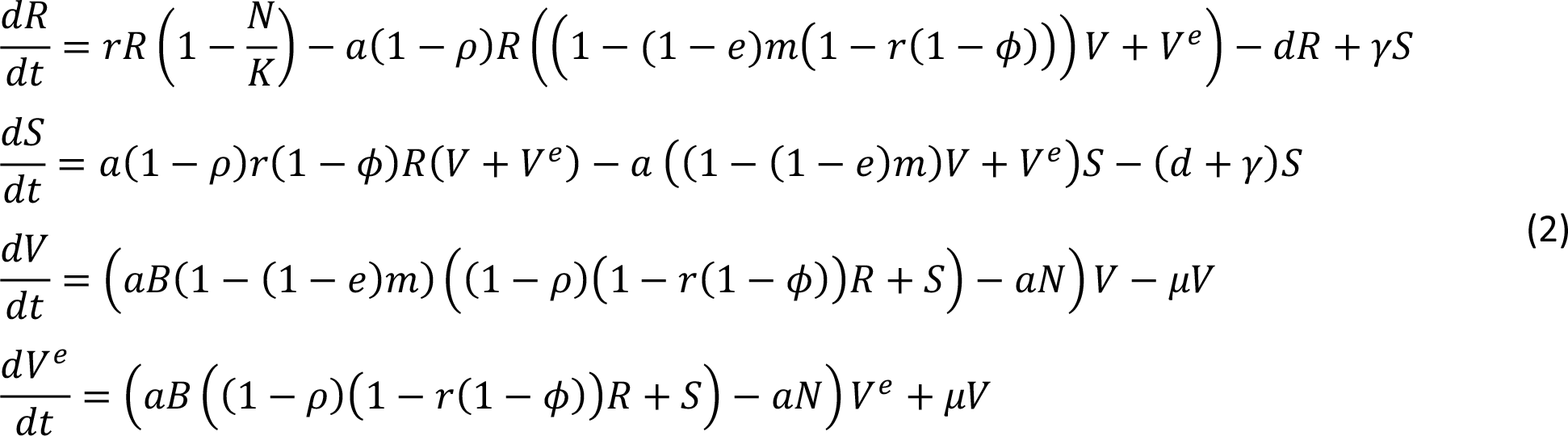

We use this model to explore what happens if we allow some mutation to occur between *V* and *V^e^* (i.e., *μ* = 0.0001), showing that the tipping point for phage amplification shifts to higher initial phage densities as the strength of CRISPR immunity increases (see Fig 4b).

Another way to formalise the evolution of escape mutation (focusing on the frequency *f^e^* of the mutant) where *f^e^* is the frequency of the mutated virus:

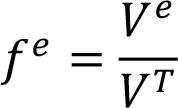

with *V^T^* = *V^e^* + *V* and:

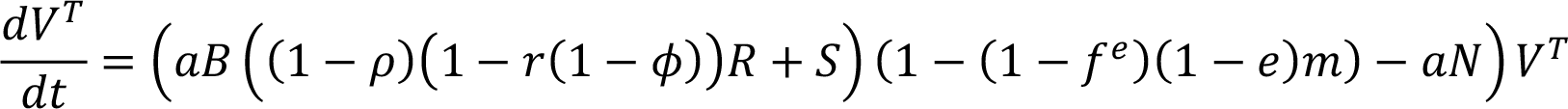

In other words, the phage population *V^T^* can only grow when:

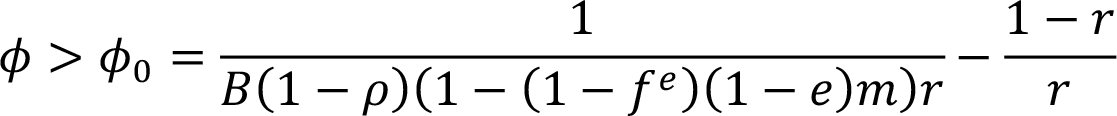

Yet, if one introduces a large density of phages in the host population they will immunosuppress a fraction *S*⁄*N* of the cells. This will yield the following threshold

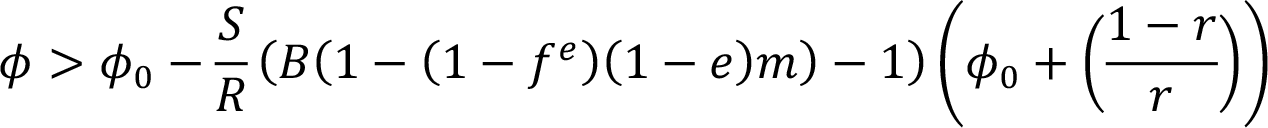

The change in mutant frequency is:

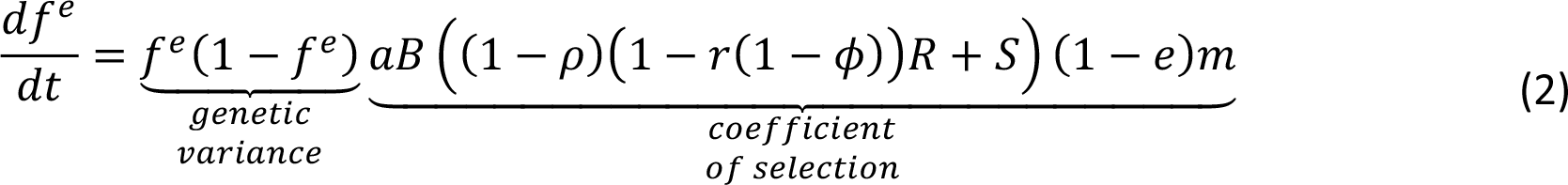

The equation (2) captures what parameters govern the speed at which the mutant virus is expected to increase in frequency. In particular, higher *m* (stronger MADS), higher *ϕ* (stronger Acr), lower *ρ* or *r* (less effective CRISPR resistance, i.e., lower numbers of spacers in the CRISPR array) promote the evolution of the mutant virus. Besides, the parameter *γ* may also affect the strength of selection via its effect on the quantity of *R* cells. When *γ* is large, the immunosuppressed cells recover their immunity very fast, the density *R* increases which favors the increase in *V^e^* frequency.

## Data availability

Source data associated with main figures and Extended Data Figures will be provided prior publication. Ordinary differential equations generated for mathematical modelling are included in Methods. Sequencing data have been deposited in NCBI under the BioProject accession number PRJNA905210.

## Code availability

Bacterial and archaeal RefSeq assemblies and scripts used to carry out bioinformatic analyses are publicly available as indicated in Methods and Supplementary Tables.

