## Supplementary_Notes for "Bacterial defences interact synergistically by disrupting phage cooperation"

### A novel defence system coined MADS

To obtain insight on the putative mechanism of MADS, we first analysed whether the genes forming the *mad1-8* operon carry domains with potential defence-related functions. Several domains could be identified, some of which are also found in other defence systems (Fig. 1b). The gene *mad1* encodes a helix-turn-helix domain protein with homology to Redirecting phage packaging protein C (RppC; PDBe: 6HN7; 92.8% confidence, Phyre2), a protein involved in packaging of phage-inducible chromosomal islands^1^. The gene *mad2* is predicted to encode an N6-Methyltransferase across methodologies. Gene *mad3* encodes a putative AbiEii toxin (Pfam-A prediction), and contains a predicted DNA-binding domain and a putative C-terminal Toprim domain that is, amongst others, found in bacterial and archaeal nucleases of the OLD family (various HHpred matches). The Mad4 protein contains a domain homologous to DUF4276 (99.78% probability, HHpred). All tools used predicted that Mad5 is a Restriction-Modification Specificity subunit and *mad6* encodes a protein kinase. Mad7 has homology to a recently-released structure of DNA phosphorothioation (PT)-dependent restriction protein DndG (PDBe: 7EXX, 99.53% probability, HHpred), and Mad8 has homology to HerA (PDBe: 42DI, 100% confidence, Phyre2), a DNA double stranded break repair helicase. While potential mechanisms for how infections are being sensed by MADS and how resistance is being mediated are not immediately obvious from this analysis, we predict, based on the methylase and specificity domains of Mad2 and Mad5, that MADS uses methylation-based self/non-self discrimination – a mechanism common to many bacterial innate immune systems, such as Restriction-Modification, BREX and DISARM^2^.

**A genomic island contains putative new defence systems, including a MADS-like system**

Amongst the 358 genomic islands that contain at least one defence system, 55% (196/358) carries genes belonging to the ‘DMS_other’ category, which recognises fragmented and non-assigned DNA-based defences (Simon Jackson, personal communication), suggesting the possibility of the existence of novel or incomplete defence systems in this location.

To further annotate the ‘DMS_other’ annotations on the genomic islands, DefenseFinder^3^ was used and found additional defence systems in 73% of the islands containing DMS_other (143/196), with RM Type IV, PsyrTA and Shango being among the most common systems to be identified (Extended Data Table 3). However, DefenseFinder was not able to find additional defence systems in the remaining 53 islands. This therefore suggests that these defence islands likely contain further novel defence systems. Manual inspection of these islands revealed one potential novel defence system that is highly similar to MADS, which we therefore called MADS-like (Fig. 2b and Extended Data Fig. 4b).

In brief, both systems encode a N6-DNA methylase, namely Mad2 and Mad-like3 (Madl3), a restriction endonuclease specificity subunit (Mad5 and Madl4) and a protein kinase (Mad6 and Madl11). Both systems carry genes at the beginning of the operon (*mad1* and *madl1*) encoding predicted DNA-binding HTH domains which may act as regulators. Notably, Madl1 encodes a WYL domain which have been shown to regulate some phage defence systems^4^. In addition, predicted protein domains for Madl9 include helicase/translocase which is also found in Mad8. Madl6 and Mad3 both encode predicted ATPase domains although HHpred hits for these two proteins are dissimilar. In contrast, MADS encodes a DUF4276 domain (Mad4) and a phosphorothioation-dependent restriction protein DptG (Mad7) that are not found in MADS-like, although the C-ter of Madl9 also contains a predicted restriction protein DptH domain which overlaps the helicase, translocase domain. Moreover, MADS-like encode four hypothetical proteins for which no significant prediction could be found.

Interestingly, defence islands carrying MADS-like were recently identified in *P. aeruginosa*^5^. These islands carry a conjugation system and plasmid partitioning proteins^5^ instead of a number of MGE- and mobility-associated genes (e.g., phage integrase, phage protein, insertion sequences) (Extended Data Fig.4b).

Apart from MADS-like, these islands also encode Type IV RM and Tiamat defence systems, DUF932 and DUF2357 domain-containing proteins, a site-specific integrase and a YqaJ recombinase, amongst others (Extended Data Fig.4b). Collectively, these analyses identify a hotspot in the *P. aeruginosa* genome for innate defence systems that may interact with the adaptive CRISPR-Cas system to determine the outcome of a phage infections.
