## Extended_Data for "Bacterial defences interact synergistically by disrupting phage cooperation"

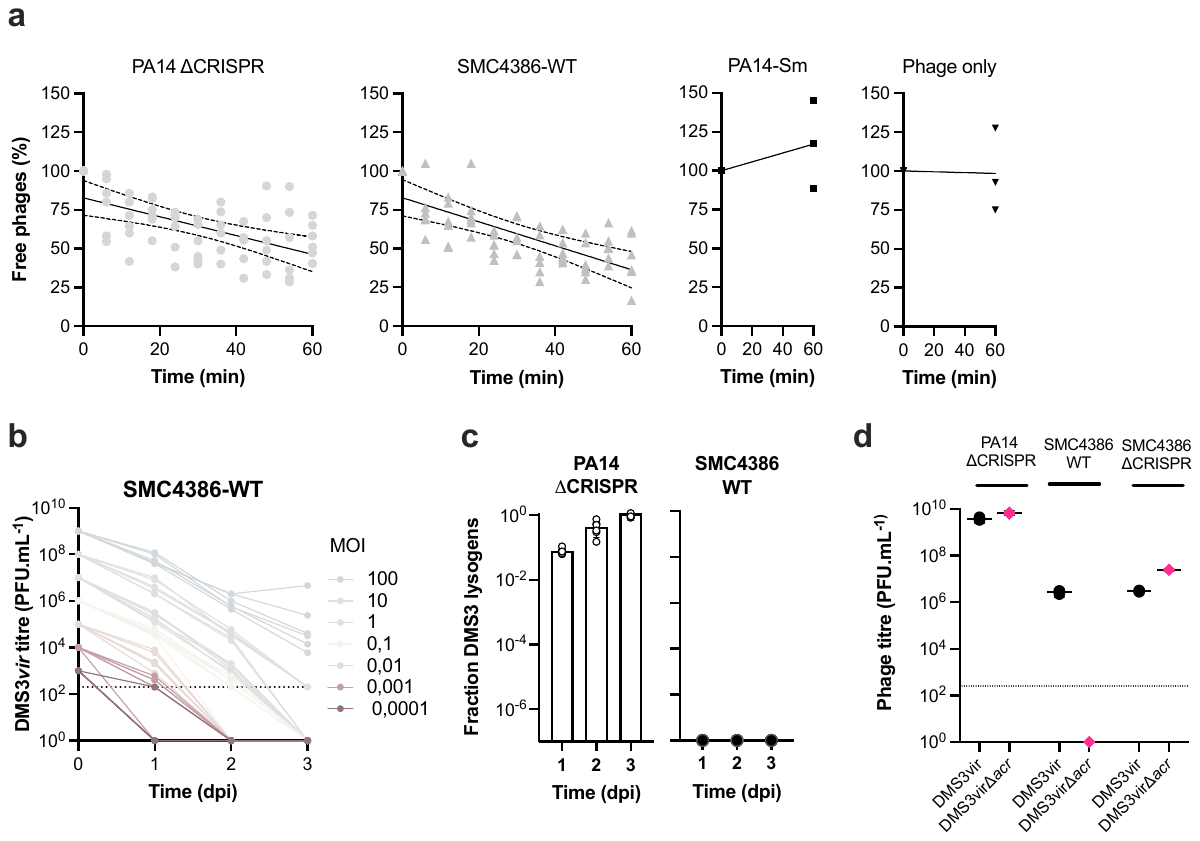

**Extended Data Fig.1 | Phage DMS3 adsorbs to *P. aeruginosa* SMC4386 but does not successfully infect this CRISPR immune strain despite a functional *acr* gene. a**, Adsorption curves of phage DMS3*vir* on strains PA14∆CRISPR (positive control), SMC4386, PA14 surface mutant (Sm) that lost phage receptor (negative control). The “phage only” panel shows the percentage of free phage detected in a flask incubated without bacteria. Individual data are shown for 6 or 3 independent replicates. Lines show linear regressions. **b**, Titre of phage DMS3*vir* over 3 days upon infection of strain SMC4386-WT with indicated initial multiplicity of infection (MOI). Individual data are shown for 6 biologically independent replicates. Horizontal dotted line shows limit of detection. **c**, Fraction of the bacterial population carrying the DMS3 prophage (lysogens) in PA14∆CRISPR or SMC4386-WT backgrounds. Individual and mean data are shown for 6 independent replicates. **d**, Titre of DMS3*vir* or DMS3*vir*∆*acr* measured on PA14∆CRISPR, SMC4386-WT or SMC4386∆CRISPR. Individual and mean data are shown for 3 independent replicates. Horizontal dotted line shows limit of detection.

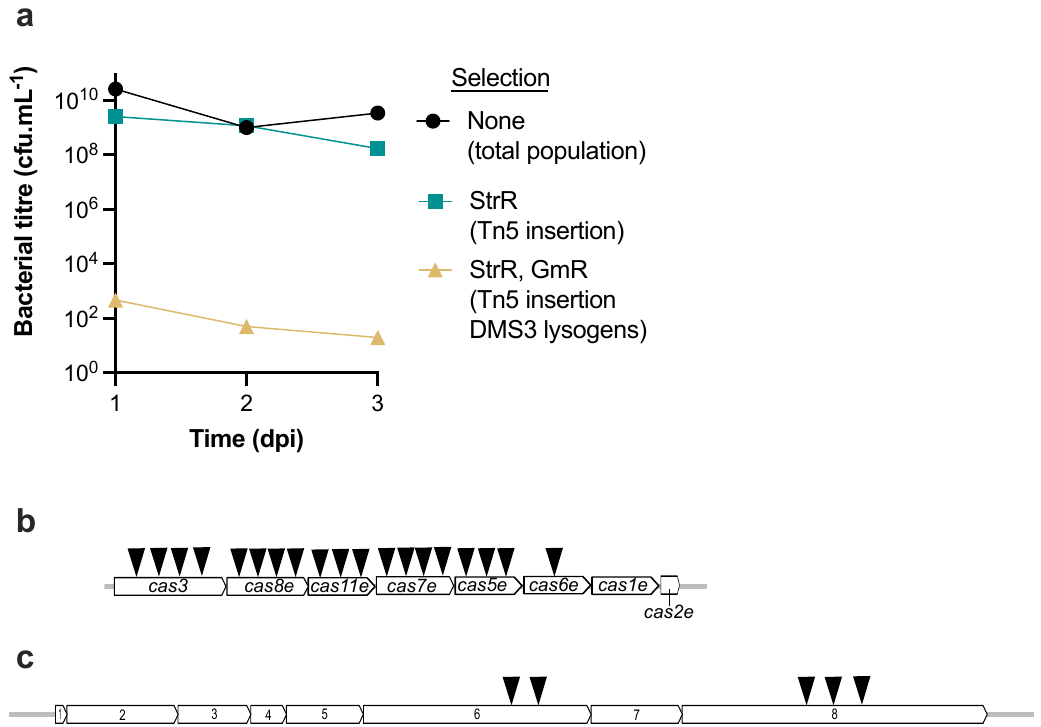

**Extended Data Fig.2 | Identification of genes involved in resistance against DMS3 phage infection**. **a**, A population of SMC4386-WT strain subjected to transposon mutagenesis (Tn5, carrying streptomycin resistance gene, StrR) was infected by a mutant phage DMS3 carrying a gentamycin resistance gene (GmR). The density of bacterial population was followed for 3 days post infection (dpi) and plated either on non-selective (LB) media (total population), on LB with streptomycin (bacterial clones with Tn5 insertion) or on LB with both streptomycin and gentamycin (bacterial clones with both Tn5 and DMS3 insertion). **b**, Tn5 insertions in the CRISPR locus upon mutagenesis of SMC4386-WT. The number of triangles indicates how many times Tn5 insertions were identified in corresponding genes (e.g.: 4 different clones have the Tn5 inserted in the *cas3* gene, 1 clone has the Tn5 inserted in the *cas6* gene). **c**, Tn5 insertion in the MADS locus upon mutagenesis of either the SMC4386-WT strain or the isogenic ∆CRISPR strain.

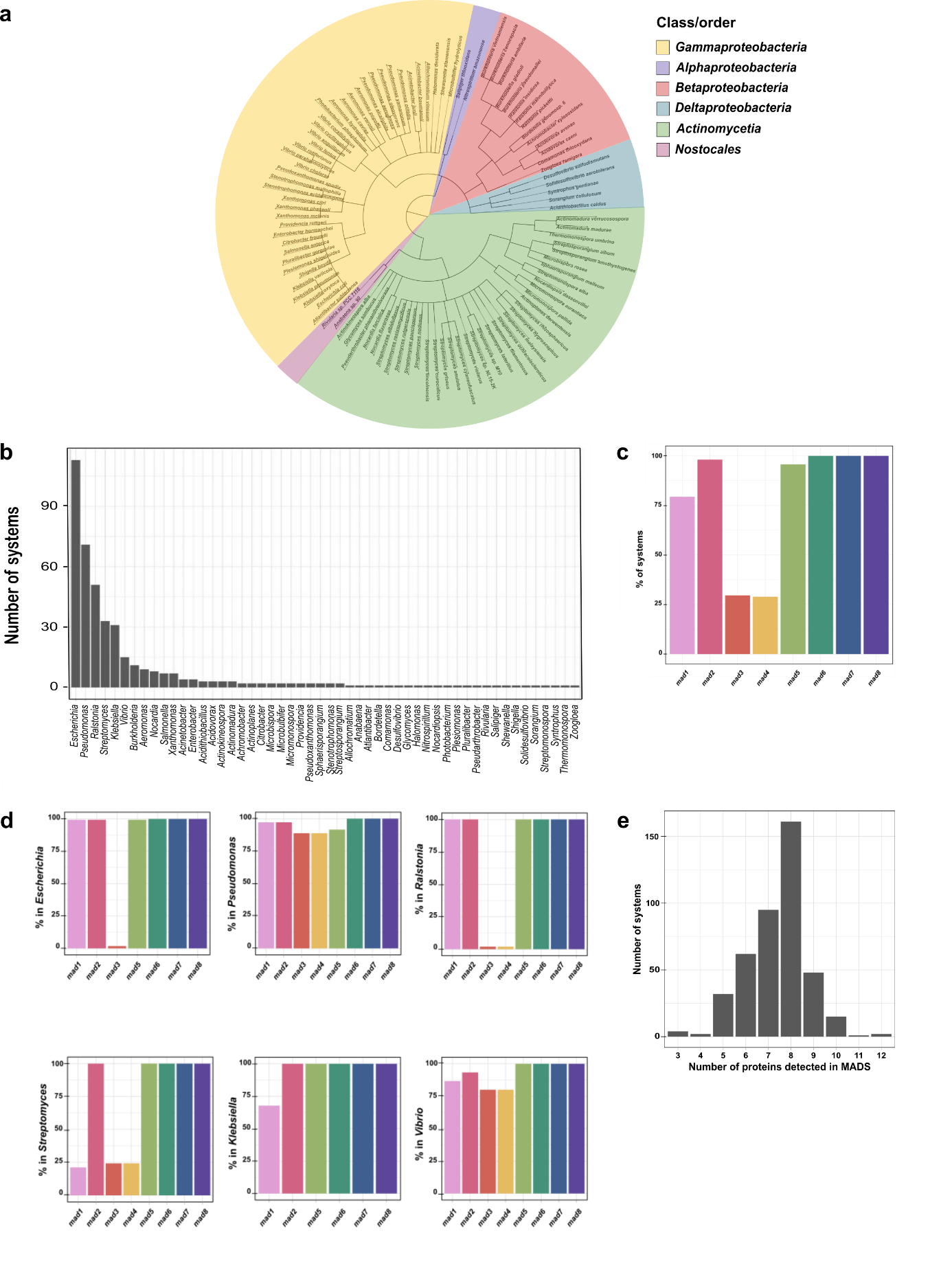

**Extended Data Fig.3 | Distribution and configuration of MADS loci in bacterial genomes. a**, Minimal tree showing the taxonomic distribution of MADS across bacterial phylogeny. **b**, The number of MADS detected in each genus in descending order. **c-d**, Percentage of systems containing each *mad*gene (*mad1-8*) within the detected MADS across all RefSeq genomes (n=422) (**c**) and percentage of systems containing each *mad* gene (*mad1-8*) for each genus where 15 or more systems were identified (**d**). **e**, The number of systems detected according to total number of proteins (>8 is possible where there are multiple hits to the same protein HMM, hidden Markov model).

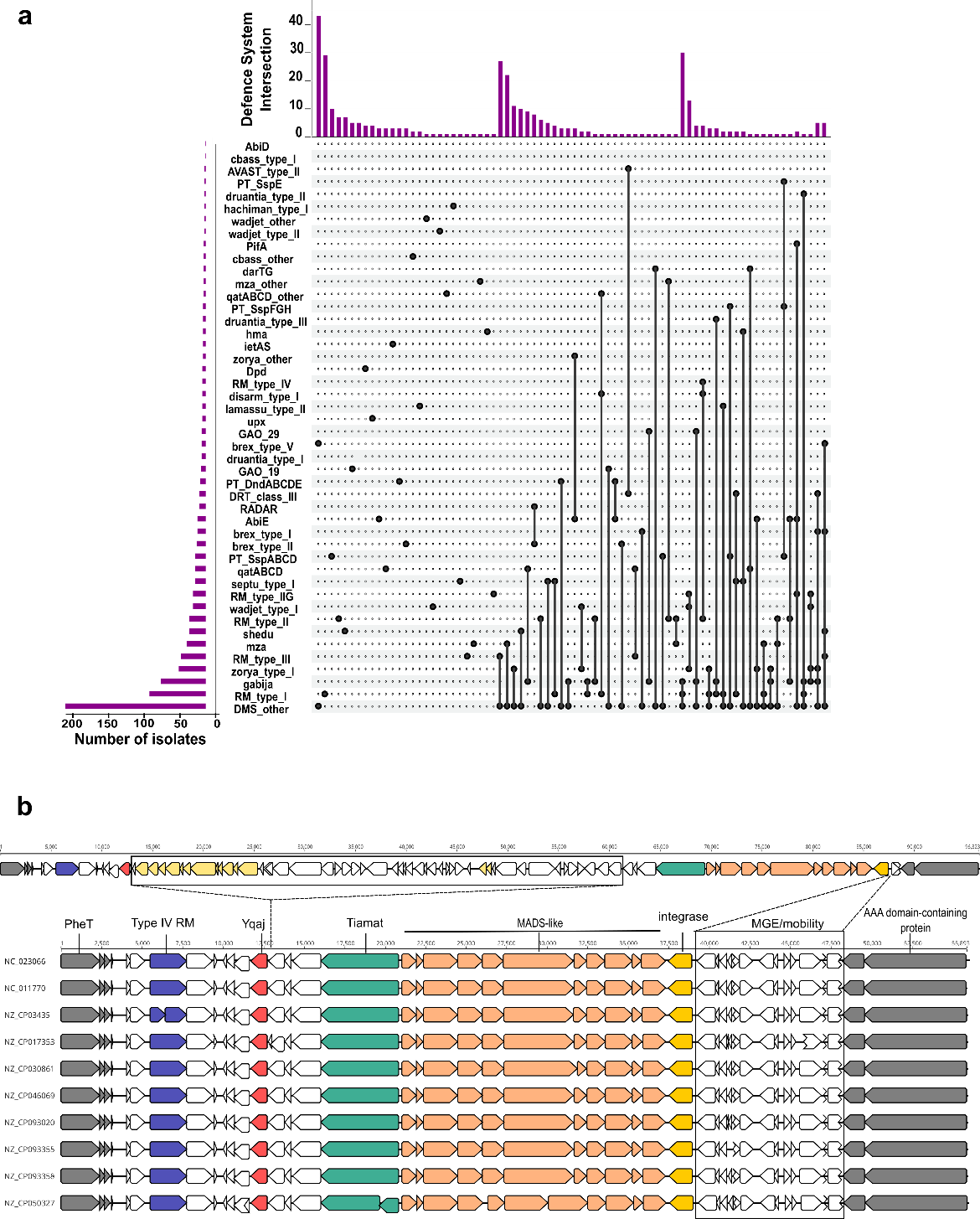

**Extended Data Fig.4 | Known and potential novel defence systems. a,** UpSet plot indicating the co-occurrence of defence systems (identified with PADLOC) in genomic island. Genotypic defence system profiles are shown in the combination matrix in the centre panel, where each column represents a unique genotypic presence/absence profile. Each black dot represents the presence of a defence system. The vertical bar plot above the matrix displays the number of isolates with a genotypic profile. The horizontal bar plot on the left of the matrix illustrates the proportion of isolates that contain each defence system. **b,** Ten defence islands that carry the MADS-like system (orange) have nearly identical genetic architecture. Blue, red, green and yellow arrows highlight Type IV RM, Yqaj recombinase, Tiamat and site-specific integrase, respectively. Grey arrows mark the boundaries of the defence islands. The top panel indicates an additional defence island carrying the MADS-like system identified in Johnson *et al* (ref ^40^). Boxes show groups of genes that are present in the top island but not in the bottom island, and vice versa. Light yellow arrows in the top box highlight genes encoding conjugation system and plasmid partitioning proteins.

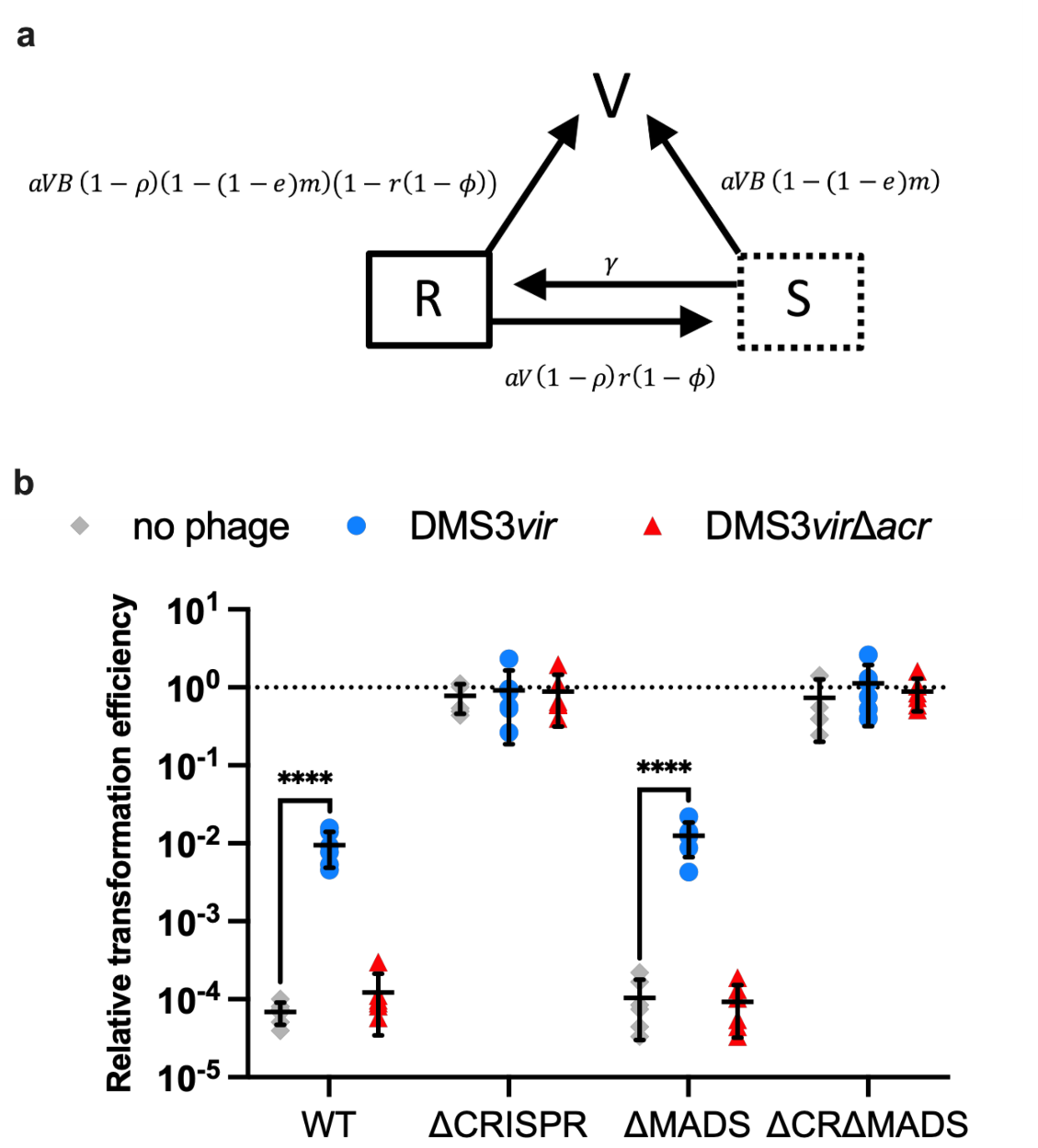

**Extended Data Fig.5 | AcrIE3 induces CRISPR immunosuppression in *P. aeruginosa* SMC4386 carrying MADS. a,** Lifecycle of Acr-phages on host that carries both CRISPR-Cas and MADS (see Methods for a detailed description of the model). We consider a multi-resistant bacterium $R$ which carries both CRISPR-Cas and MADS. Exposition to Acr-phages leads to a new class of immunosuppressed cells $S$ for which CRISPR resistance is impaired (but activity of MADS is maintained). The efficacy of CRISPR resistance is noted $\rho$ (for the ability to block infection) and $r$ (for the ability to induce immunosuppression after being infected by an Acr-phage). The efficacy of MADS is noted $m$ and only blocks infection (the completion of lytic infections). The efficacy of the Acr protein is noted $\phi$ and the efficacy of escape mutations against MADS is noted $e$. **b**, Transformation efficiency of a CRISPR-targeted plasmid relative to a non-targeted plasmid into SMC4386-WT and mutants lacking either CRISPR-Cas, MADS or both, following pre-infection with either no phage, phage DMS3*vir* (which encodes *acrIE3*) or DMS3*vir*∆*acrIE3*. See Methods for explanations on the calculation of relative transformation efficiency. Individual and mean data is shown for 6 biologically independent replicates. Errors bars show standard deviation. **** indicates p<0,0001, (two-way Anova – Dunnett’s test).

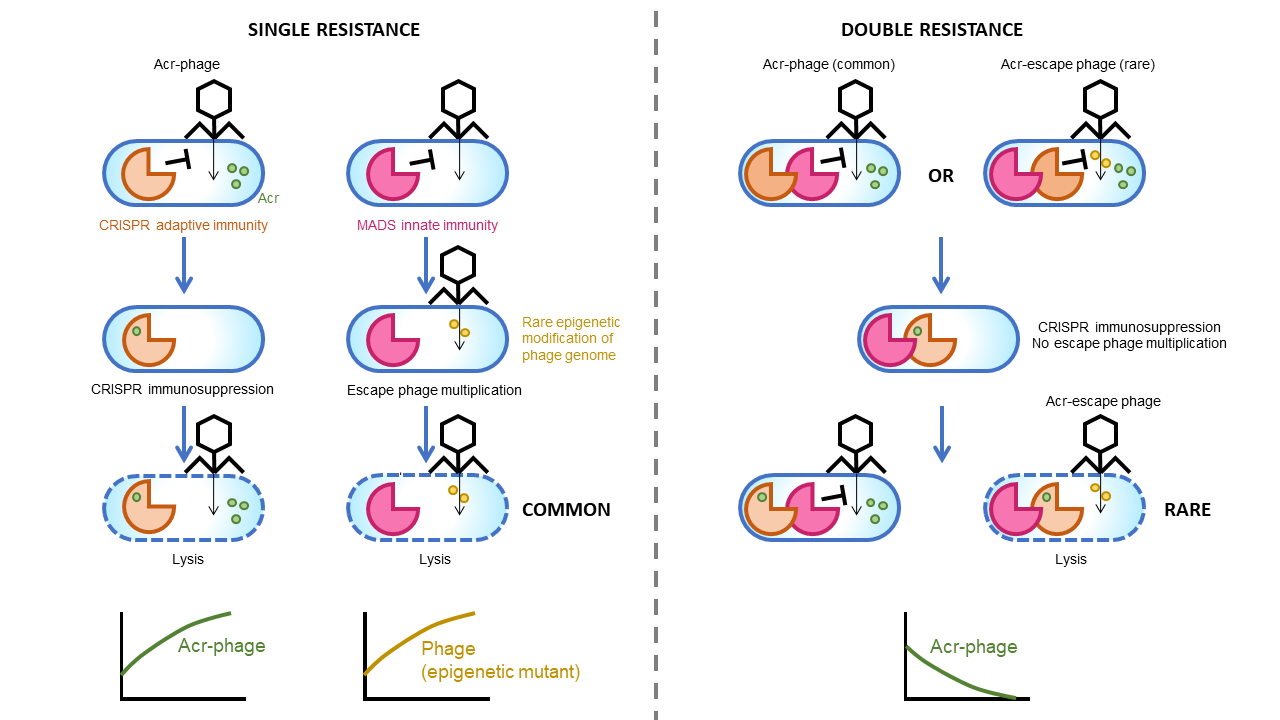

**Extended Data Fig.6 | Synergy between CRISPR-Cas and MADS.**

Schematisation of different phage epidemiological outcomes based on the interaction between distinct types of bacterial resistance (single and double) and the deployment of phage counter-defence strategies (Acr-genes and methylation of phage genome).
